# Downregulation of a satiety signal from peripheral fat bodies improves visual attention while reducing sleep need in *Drosophila*

**DOI:** 10.1101/808998

**Authors:** Deniz Ertekin, Leonie Kirszenblat, Richard Faville, Bruno van Swinderen

## Abstract

Sleep is vital for survival. Yet, under environmentally challenging conditions such as starvation, animals suppress their need for sleep. Interestingly, starvation-induced sleep loss does not evoke a subsequent sleep rebound. Little is known about how starvation-induced sleep deprivation differs from other types of sleep loss, or why some sleep functions become dispensable during starvation. Here we demonstrate that downregulation of unpaired-2 (*upd2*, the *Drosophila* ortholog of leptin), is sufficient to mimic a starved-like state in flies. We use this ‘genetically starved’ state to investigate the consequences of a starvation signal on visual attention and sleep in otherwise well-fed flies, thereby sidestepping the negative side-effects of undernourishment. We find that knockdown of *upd2* in the fat body is sufficient to suppress sleep while also increasing selective visual attention and promoting night-time feeding. Further, we show that this peripheral signal is integrated in the fly brain via insulin-expressing cells. Together, these findings identify a role for peripheral tissue-to-brain interactions in the simultaneous regulation of sleep and attention, to potentially promote adaptive behaviors necessary for survival in hungry animals.

**Author Summary:** Sleep is important for maintaining both physiological (e.g., metabolic, immunological, and developmental) and cognitive processes, such as selective attention. Under nutritionally impoverished conditions, animals suppress sleep and increase foraging to locate food. Yet it is currently unknown how an animal is able to maintain well-tuned cognitive processes, despite being sleep deprived. Here we investigate this question by studying flies that have been genetically engineered to lack a satiety signal, and find that signaling from fat bodies in the periphery to insulin-expressing cells in the brain simultaneously regulates sleep need and attention-like processes.

## Introduction

Behavioral decisions in animals are formed by integrating internal states with external stimuli and prior experience. The need for sleep and food are two such internal states, and satisfying both of these homeostatic processes seems equally important for survival. Yet sleeping and feeding are also mutually exclusive: they cannot happen at the same time. Under environmentally challenging conditions, mutually exclusive behaviors therefore need to be prioritized in order to maximize survival. Behavioral performance depends as much on getting a good night’s sleep as on being well-fed.

Both sleep and feeding regulation have been extensively studied in different animal models, as well as in humans [1–7]. Yet, how their pathways intersect and influence each other remains unclear. Given the alarming increase in the number of people suffering from both sleep and metabolic disorders [8, 9], to understand how these two processes interact at the level of neural circuits and molecular pathways is of significant interest. *Drosophila melanogaster* has been a pivotal model system to study both sleep and feeding regulation [10, 11]. Sleep in flies has been shown to fulfill key criteria for identifying sleep in other animals, such as increased arousal thresholds and homeostatic regulation [12, 13], so the fly is a promising avenue for understanding sleep and feeding regulation at a circuit level. Additionally, cognitive readouts such as visual attention paradigms are increasingly available for *Drosophila* research [14–16], providing relevant functional assessments of manipulations that could impact sleep and feeding.

Generally, the effect of feeding on sleep has been studied by altering dietary components, or by more severe interventions such as starvation [17–20]. However, studying this relationship via nutritional manipulations introduces numerous secondary factors (e.g., metabolic processes or energy levels), confounding any analysis of potential interactions between satiety/starvation signals and sleep processes. We therefore decided to utilize a genetic strategy in *Drosophila* to downregulate a satiety signal, Unpaired-2 (*upd2*), thereby mimicking a ‘starved’ state in flies, allowing us to assess the effect of genetic starvation on sleep and attention simultaneously. *upd2* is a functional ortholog of vertebrate leptin, which has similar structure to type-I cytokines [21, 22]. Similar to leptin, secretion of *upd2* is dependent on nutrient intake [21, 23], and is secreted from the fly counterpart of adipocytes, or fat bodies.

Starvation has several consequences on the behavior of animals, with one of the most striking ones being suppression of sleep [17]. Normally, sleep deprivation in flies and other animals leads to an increase in sleep drive and a homeostatic sleep rebound [12,13,24], as well as impaired cognitive capacities such as visual learning [25, 26] and attention [27]. Yet, starvation-induced sleep loss seems to absolve animals from sleep need and some of the functional consequences of sleep loss [27–29]. The mechanisms supporting this surprising effect are unclear, and it is unknown what aspects of cognition are preserved under this regime. We used *upd2* mutants and tissue-specific *upd2* knockdown to address the consequences of a prolonged starved-like state. We demonstrate that lack of a satiety signal disrupts day-time sleep and leads to night-time hyperphagia (increased feeding at night). While sleep deprivation typically impairs attention, we found that genetically-starved animals had improved attention, even though they slept less. Finally, we show that *upd2* regulates sleep and attention via the *insulin-like peptide 2* signaling in the brain. Our results highlight a role for peripheral signaling in co-regulating cognition and sleep as a function of nutrition.

## Results

### *Upd2* mutants have irregular feeding and fragmented sleep

Homozygous *upd2* deletion mutants (*upd2^Δ^)*, which lack the 5’UTR and the first 89 amino acids of the protein [30], have been shown to be smaller and slimmer than control animals [21]. We first measured the food intake of mutant animals, to address if the difference in their body size (Figure 1A) was due to a decrease in feeding. We used an optimized version of the capillary feeding (Café) assay [31], and tracked their feeding over 24 hours. In agreement with previous findings, there was no significant change in total food consumption in *upd2^Δ^* mutants, compared to controls (Figure 1B) [21, 32]. However, when we looked at day and night feeding separately, we noticed that the mutants were mostly feeding during the night, which was opposite to the feeding rhythm of the background controls (Figure 1C). The difference in feeding timing was independent of light entrainment, excluding the possibility of a circadian influence (Figure S1A). This suggests that *upd2^Δ^* mutants are feeding when they normally should be achieving most of their deeper sleep [33].

**Fig 1.**
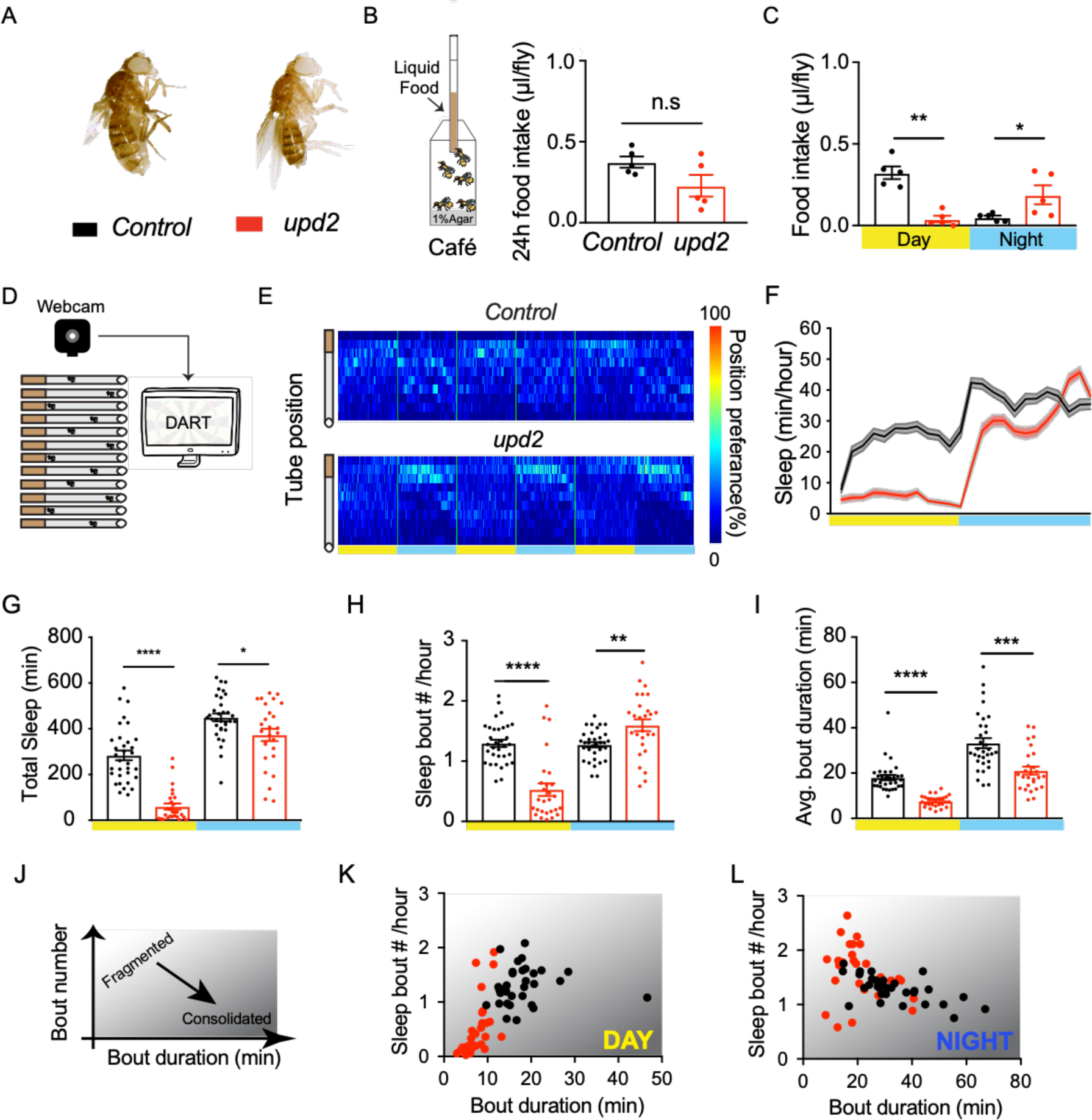
*Upd2* mutants have altered feeding behavior and fragmented sleep. (A) Homozygous *Upd2* deletion mutant females (right, red) are slimmer and smaller in size compared to their background controls (left, black) (B) Food intake was measured with Café assay, with 5 flies/chamber over 24 hours (n=40 per genotype). Bottom of the chamber had 1% Agar to prevent desiccation. Liquid food (5%Sucrose/Water) was presented in a microcapillary. Total food consumption of *upd2* mutant flies (red) was similar to their background control (black). (C) Mutants had lower consumption during day-time. However, they had a significant increase in their night-time feeding. (D) *Drosophila* ARousal Tracking (DART) was used to measure sleep duration in *upd2* mutants. 3-5-day old female virgins were placed in glass tubes and sleep was tracked over 3 days. (n=31-32, per genotype). (E) Average position preference heatmaps show that *upd2* mutants have an increased presence at the food side at night (blue bars), whereas controls remain in the center. (F) 24hr sleep profile of *Upd2* mutants compared to control. Yellow and blue bars represent light and dark periods, respectively. (G) *upd2* mutants had a significant reduction in average sleep duration for both day and night. (H) The number of sleep bouts was reduced during day-time and increased during night-time. (I) The average bout duration of *upd2* mutants was reduced for both day and night. (J) Bout number plotted against bout duration is reflective of sleep quality. Sleep is more fragmented when bout durations are short and bout numbers are high, whereas sleep is consolidated with low bout numbers and longer bout durations. (K-L) Total bout number plotted against average bout duration (min) showed that *upd2* mutants had fragmented day and night sleep. *Student’s t-test for normally distributed data or Mann-Whitney U rank-sum test for nonparametric data was used to compare datasets. *P <0.05, **P <0.01, ***P <0.001, Error bars show SEM*.

Nutritional state has been shown to influence sleep duration, as well as quality of sleep [17, 34]. We used the *Drosophila* Arousal Tracking (DART) system [33] to monitor sleep duration in *upd2^Δ^* (Figure 1D). In accordance with our Café results (Figure 1C), position preferences of flies in tubes revealed that *upd2* mutants consistently stayed near the food during the night, unlike the background controls (Figure 1E). *Upd2^Δ^* displayed a regular day-night sleep profile (sleeping less during the day and more at night, Figure 1F), however they require significantly less sleep than control flies, especially during the day (Figure 1G). Closer analysis revealed a decreased number of sleep bouts, which were shorter in duration during the day (Figure 1H-I). During night-time however, mutants had increased sleep bout numbers, which were shorter in duration (Figure 1H-I). These results also indicate that sleep in *upd2* mutants is more fragmented than in control animals, both during both day and night (Figure 1J-L). Overall, the observed decrease in sleep duration, sleep quality, and the mis-regulation of feeding suggested a persistent ‘starved-like’ state in these mutants, presumably resulting from the absence of a satiety signal that normally results from adequate feeding. However, rather than feeding or sleeping during the day, the mutant flies are mostly walking persistently back and forth in their chambers with constant speed (Figure S2).

### *upd2* secretion from fat bodies regulates sleep quality

The *Drosophila* fat body is the main tissue for energy storage, fulfilling functions similar to the mammalian adipose tissue and liver [35, 36]. In *Drosophila*, Upd2 is mainly secreted from the fat body [30]. To test the role of adult *Upd2* secretion on sleep and feeding, we used RNA interference (RNAi), expressed via an adult fat-body (FB) specific driver line (Yolk-Gal4) [37]. We measured food consumption in flies where *upd2* had been downregulated in the fat body specifically, to determine if this recapitulated the effects seen in *upd2* mutants. This is indeed what we found: when we compared day versus night, we found that day-time feeding remained similar to controls, while night-time feeding increased (hyperphagia) (Figure 2A), similar to what we observed in *upd2* mutants (Figure 1C). Importantly, this shows that *upd2*-downregulated flies are not undernourished, compared to controls. Also similar to the *upd2* mutant phenotype, fat-body specific *upd2* knockdown resulted in a significant suppression of day-time sleep (Figure 2B-C). Together with the preceding results, this suggests that the satiety signal affecting sleep and feeding emanates from the fat body.

**Fig 2.**
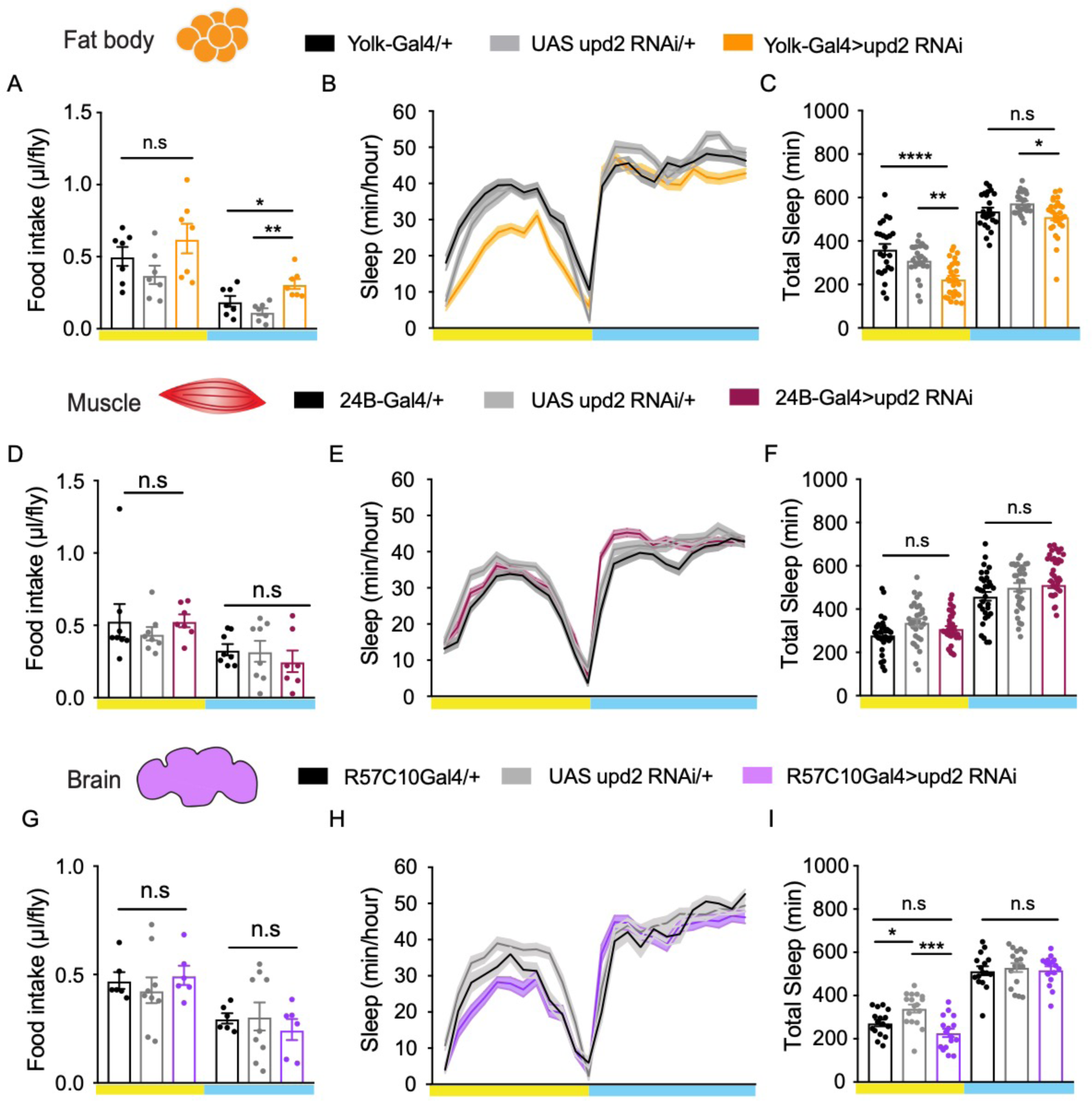
Upd2 expression in the fat body is required for sleep and feeding regulation. (A) Day-time feeding (left panel-yellow) in flies with fat body upd2 knockdown (orange) was similar to genetic controls; *Yolk-Gal4/+ (black)* and vs *UAS-Upd2^RNAi^/*+ (grey). Night-time feeding (right panel-blue) was significantly reduced in knockdown flies. (B) 24h sleep profile of flies with *upd2* knockdown (orange line) compared to parental controls (black, *Yolk-GAL4/+*; grey, *UAS-Upd2^RNAi^/+*) (mean± SEM, n=24-28 per genotype) (C) Sleep duration in knockdown flies was significantly reduced during the day against both controls. Night-time sleep was only reduced compared to *UAS-Upd2^RNAi^/* +. (D) Muscle specific knockdown of *upd2* by using *24B-Gal4 (maroon)* did not alter day or night-time food intake (right) (n=20) or (E-F) sleep duration. (G) Knockdown of *upd2* pan-neuronally using *R57C10-Gal4* (purple) also had no effects on food intake (n=20) or (H-I) on sleep (n=16-17). *One-way Anova (with Tukey’s post-hoc test) for normally distributed data or Kruskal-Wallis test (with Dunn’s multiple comparison test) for nonparametric data was used to compare datasets. *P <0.05, **P <0.01, ***P <0.001, Error bars show SEM.* Yellow and blue bars indicate day and night, respectively.

*Upd2* is also found to be expressed in muscle tissue [30, 38], so it remained possible that the satiety signal was not localized to the fat body. To determine whether the effect on sleep and food consumption was specific to expression in the fat body, we assessed the effect of *upd2* knockdown in muscle tissue. For this, we employed a muscle-specific driver, 24B-Gal4 [39]. We found that muscle-specific knockdown of *upd2* had no effect on food consumption or sleep behavior (Figure 2D-F). We then asked whether downregulation of *upd2* in neuronal tissue could lead to a starved-like state. One of the other *unpaired* ligands, *upd1*, is expressed in a small cluster of cells in the brain [32], but it is currently unknown if *upd2* is expressed in neurons. However, pan-neuronal knockdown (*via* R57C10-GAL4 [40]) of *upd2* had no effect on feeding behavior, and any effects on sleep were inconsistent compared to controls (Figure 2G-I). These results support the conclusion that the most robust effects of *upd2* on sleep and feeding result from its secretion from the fat body. We therefore subsequently focused on *upd2* signaling from the fat body.

Decreased sleep duration does not necessarily imply decreased sleep quality; flies could be sleeping more deeply in shorter bouts, and thereby still achieving key sleep functions [33, 41]. We therefore next investigated sleep intensity in *upd2*-downregulated flies. To measure sleep intensity, we delivered a series of vibration stimuli every hour, and analyzed the proportion of sleeping flies responding to the stimulus (Figure 3A). We binned the flies into two different sleep duration groups, according to the amount of time they had been asleep before the arousal test, 1-30 min or 31-60 min (Figure 3A). To accurately assess sleep intensity, we ensured that both sleep duration groups had a similar number of arousal-probing events in all of our genetic variants (Figure 3B). We found that downregulating *upd2* in the fat body resulted in significantly lighter day-time and night-time sleep during the first 1-30min of sleep (flies were more responsive), compared to genetic controls (Figure 3C, left histograms). Interestingly, later stages of sleep (31-60 min) were not significantly different compared to both controls (Figure 3C, right histograms). This suggests that lack of a satiety signal from the fat body results in decreased sleep intensity in earlier sleep stages, in addition to decreased sleep duration. Thus, both sleep duration and sleep intensity are impacted by *upd2* downregulation.

**Figure 3.**
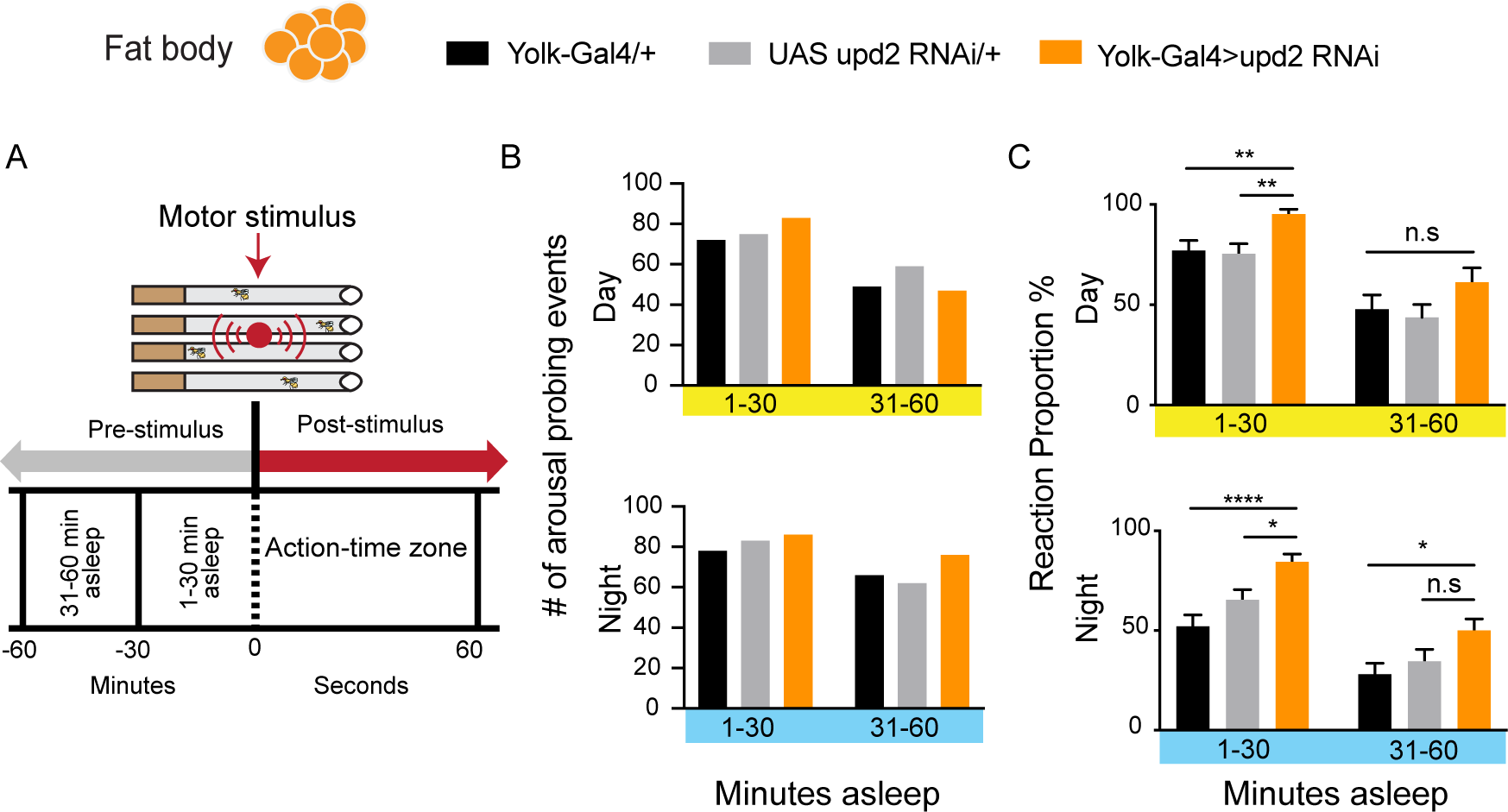
upd2-downregulation decreases sleep intensity. (A) A vibration stimulus was presented to flies, to measure sleep intensity. A stimulus train consisted of five-pulses with 0.2 s and was presented once per every hour. Flies were binned according to their pre-stimulus sleep duration (1-30 min. or 31-60 min. asleep). Reaction proportion represents the percent of immobile animals, which were responsive to the stimulus train within 60 seconds. (B) The number of animals in each bin group for both day and night were similar in both genetic controls (black and grey) and in knock down flies (orange). (C) Knockdown flies were significantly more responsive during lighter stages of sleep (1-30 min inactive) both day and night-time. Deeper stages of sleep (31-60 min inactive) were not affected. *One-way Anova (with Tukey’s post-hoc test) for normally distributed data or Kruskal-Wallis test (with Dunn’s multiple comparison test) for nonparametric data was used to compare datasets. *P <0.05, **P <0.01, ***P <0.001, Error bars show SEM.* Yellow and blue bars indicate day and night, respectively.

### Tracking sleep and feeding behavior in individual animals

In our preceding experiments, we found that *upd2* downregulation has correlated effects on sleep and feeding behavior, although these observations were made using different assays for different sets of flies. To confirm our findings, and to further our understanding of relationship between sleep and feeding, we devised a novel open-field paradigm wherein we could monitor sleep and feeding in the same animals. In this setup, individual flies were housed in circular arenas provisioned with food in the center of each chamber (Figure 4A). The protocol for sleep tracking was kept the same as in our previous experiments in small glass tubes (see Methods). Importantly, we confirmed that the sleep profile of *upd2* knockdown flies was similar in this open-field setup, with a significant sleep reduction during the day (Figure 4B-C, compared to Figure 2B-C).

**Fig 4.**
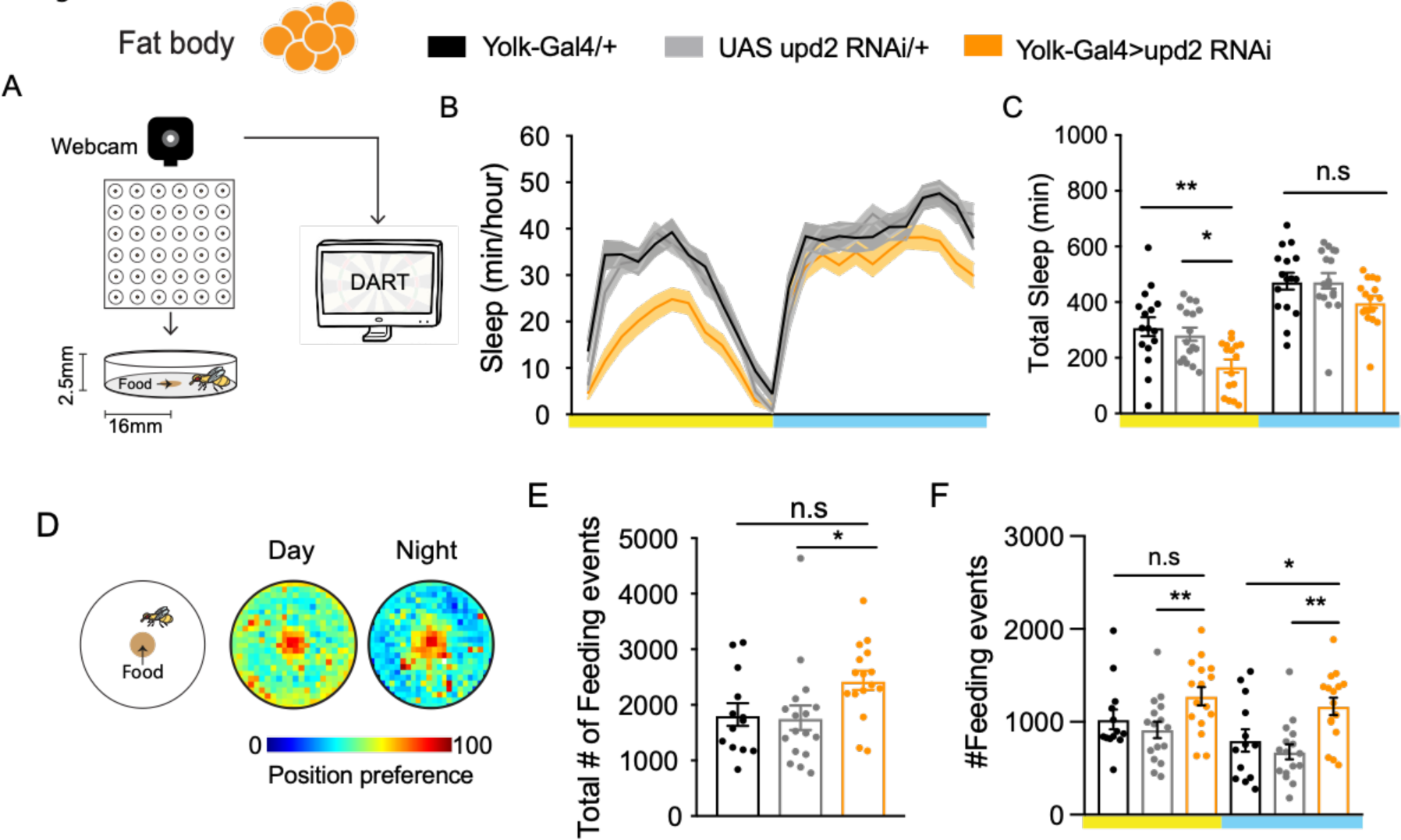
A novel open-field platform to study sleep and feeding simultaneously. (A) Schema of the open-field sleep and feeding tracking system. Flies are housed individually in arenas, with r=16mm. Each arena has a food pit in the center (R=5mm). Fly activity is monitored via a webcam and analyzed with DART. (B) 24h sleep profile of flies with *upd2* knockdown (orange line) compared to genetic controls. Sleep was tracked for 3 days (n=15-17 per genotype), (black, *Yolk-GAL4/+;* grey, *UAS-Upd2^RNAi^/+*). (C) Fat-body specific *upd2* knockdown in open-field arena showed a similar phenotype as in tubes (orange bars). (D) Exemplary average 2D-heatmap position preference plots for day and night-time, warmer colors indicate a higher probability of flies being in that position. (E) Number of feeding events for each genotype over 3 days. Total feeding events in knockdown flies was only significant compared to one of the parental controls, *UAS-Upd2^RNAi^/+*. (F) Upd2 knockdown flies had an increase in feeding events during night-time. Day-time feeding count was increased as well, was however only significantly different to one of the parental controls (*UAS-Upd2^RNAi^/+*). *One-way Anova (with Tukey’s post-hoc test) for normally distributed data or Kruskal-Wallis test (with Dunn’s multiple comparison test) for nonparametric data was used to compare datasets. *P <0.05, **P <0.01, ***P <0.001, Error bars show SEM.* Yellow and blue bars indicate light and darkness, respectively.

The circular arena setup allowed us to track the absolute location of flies in two dimensions. Heat plots revealed that flies frequently visited the food cup located in the center of the arena (Figure 4D). Similar to sleep tracking (above), feeding was tracked over multiple days (see Methods). Our automated analysis was based upon visual observation; a fly was considered to be feeding when it fulfilled 3 criteria, (1) it was located at the food, (2) its speed was less than 1mm/sec, and (3) it remained there for at least thirty seconds. We further confirmed the accuracy of the feeding tracking by preventing access to the food source: covering the food cup with parafilm decreased the number of ‘feeding’ visits to the center of the arena, whereas replacing the food with 1% agar did not (Figure S3A). This accurate estimate of feeding behavior allowed us to quantify the number of ‘feeding events’ over three days and nights in our combined paradigm (Figure 4E). When we again partitioned our analysis between day and night, we observed that downregulation of *upd2* in the fat body significantly increased the number of feeding events during the night (Figure 4F, right panel). These results are in accordance with our Café assay results (Figure 2A). This indicates that the feeding phenotype is unlikely to be an artifact of different assay conditions (e.g., the liquid food of the Café assay) or due to group housing in Café chambers. Thus, regardless of the assay employed, removing a satiety signal from the fat body reliably decreases day-time sleep and increases night-time feeding. The observed sleep reduction and sleep fragmentation resulting from *upd2* downregulation aligns with the behavior of starved animals more generally [17]. Moreover, the hyperphagia of *upd2* mutants suggests that the signal communicating that food has been consumed is not being integrated or processed, even if flies are well-fed. This starvation cue may be the cause of decreased sleep, rather than nutritional status *per se*.

### Neural correlates of starvation in the ellipsoid body R4-neurons

If *upd2* mutants are failing to process a satiety signal, then neural evidence of this genetically-induced starvation state should be evident in brain activity. We used a CaLexA (Calcium-dependent nuclear import of LexA) reporter [42] to visualize activity levels in the brains of *upd2* mutants, and compared these to starved animals. We expressed the reporter construct in the ellipsoid body R4 neurons (using R38H02-Gal4, (Figure S4A)), which have been shown to increase their activity under a starvation regime [43]. We tracked both sleep and feeding behavior in individual animals in the open-field arena (as in Figure 4) for two days, after which we dissected and imaged their brains (Figure 5A). In wild-type flies, we found that activity in the R4 neurons was significantly correlated with the time spent feeding (Figure 5B-C), but not with sleep duration (Figure S4B). Since other classes of ring neurons have nevertheless been implicated in sleep [24], we conducted additional experiments to determine if increased R4 neuron activity also affected sleep. We found that optogenetic activation of the R4 neurons (using UAS-csChrimson/R38H02-Gal4, see Methods) decreased sleep (Figure S5A-D).

**Fig 5.**
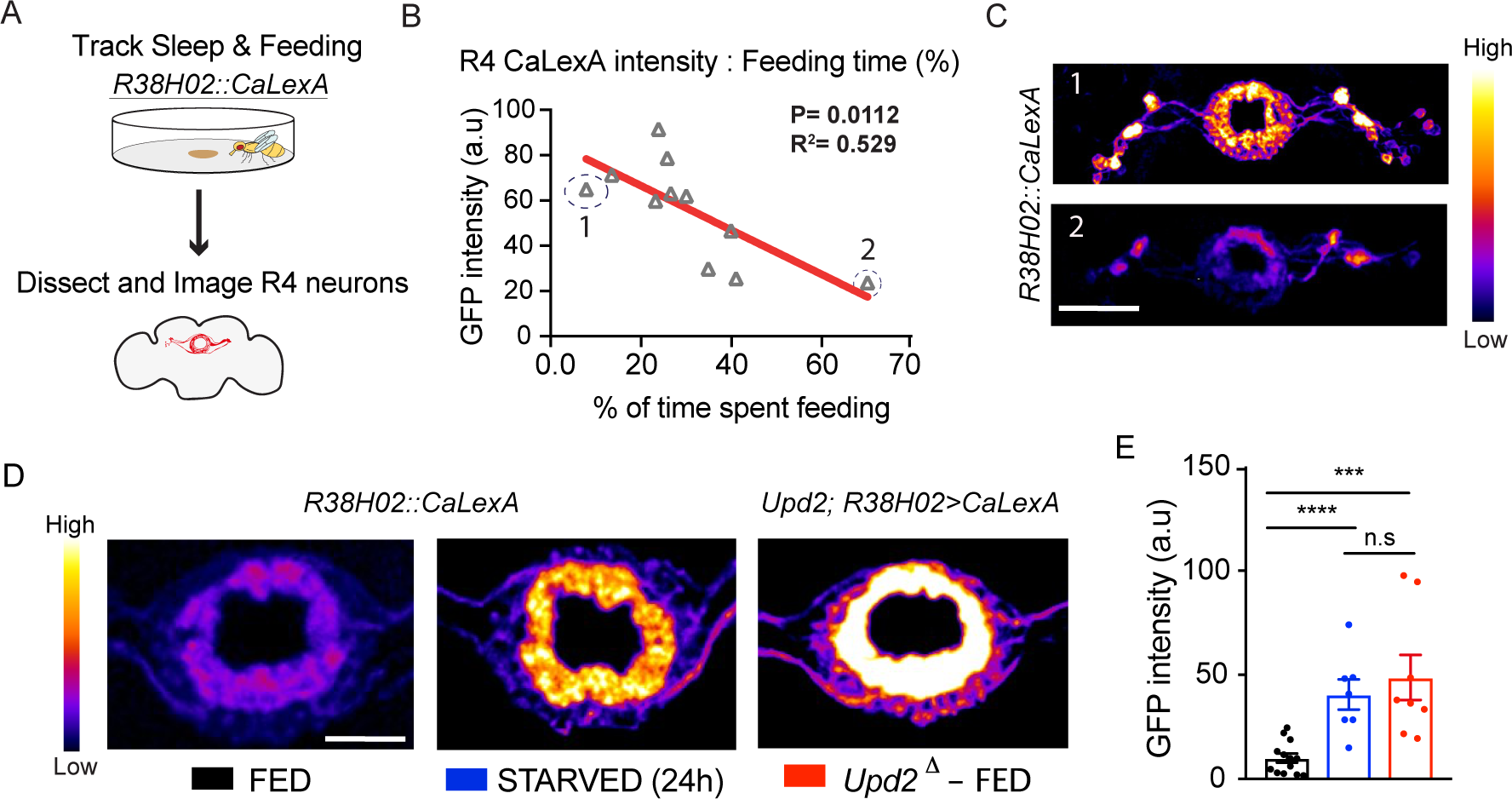
Upd2 mutants show increased/starved-like CaLexA expression in the ellipsoid body R4 neurons. (A) Flies were housed in open-field arena for 24 hours (starting at ZT0) for sleep and feeding tracking. They were then collected for dissection and brains were imaged. (B) Time spent feeding in open-field arena significantly correlated with the measured CaLexA intensity of R4 neurons. (C) Sample images presenting a high (no:1, upper panel) compared to a low CalexA signal (no:2, lower panel). n=11, Warmer colors indicate an increase in GFP intensity. (D) Representative whole-mount brain immunostaining of fed wild-type (*w^+^; R38H02-GAL4>UAS-CaLexA, left*), starved wild-type (*w^+^; R38H02-GAL4>UAS-CaLexA, center*), and *upd2* mutant (*Upd2;R38H02-GAL4>UAS-CaLexA, right*) background. Maximal intensity projections are shown in pseudo color. (Scale bar= 20µm). (E) Quantification of CaLexA signals; Starvation led to an increase in R4 activity compared to fed controls. *Upd2* mutant flies also had a significant increase in activity, which was similar to starved controls. (n=8-13 per condition). Control flies had ad libitum access to food from day 0-6. Starved group was transferred to 1% agar/starvation medium on day 5 (ZT0) until day 6 (ZT0). *For correlation analyses* two-tailed p-values for Pearson’s correlation coefficient are shown. *Student’s t-test for normally distributed data or Mann-Whitney U rank-sum test for nonparametric data was used to compare datasets. *P <0.05, **P <0.01, ***P <0.001, Error bars show SEM*.

Consistent with the above data and a previous study [43], we observed a significant increase in R4 neuron activity after 24 hours of starvation (Figure 5D, center compared to left in two example flies). We then placed the CaLexA/R38H02 reporter in an *upd2* mutant background, to investigate R4 neuron activity in these flies. Interestingly, *upd2* mutants also display increased R4 neuron activity, even though they had *ad libitum* food access and consumed similar amounts as fed controls (Figure 5D, right). The average GFP intensity of R4 neurons in the mutants was comparable to the level observed in starved wild-type flies (Figure 5E). Overall, our results are strongly indicative that lack of the *upd2* satiety signal produces a starved-like state, associated with this neural signature in the fly brain.

### A starved-like state sharpens visual selective attention

Sleep-deprived flies have been shown to have deficits in learning and memory and visual attention tasks [25,27,44]. Yet, starvation-induced sleep loss seems to preserve performance in some behavioral assays [29]. Indeed, many behavioral studies exploit starvation as a way to increase motivation and to even improve performance [45–47]. Since most sleep-monitoring assays for *Drosophila* do not provide much insight into behavioral processes beyond locomotion, it remains unclear how starvation might preserve or improve behavioral performance in spite of lost sleep.

We investigated whether a starved-like state affected visual selective attention. To study visual attention in flies, we used a modified version of Buridan’s paradigm, to track visual fixation behavior in freely-walking animals (Figure 6A) [48, 49]. To ensure that vision was normal in our genetic variants, we outcrossed them to *white^+^* so that their eye pigmentation was wild-type (see Methods). We then proceeded to test them first for simple visual behaviors, namely object fixation [50, 51] and optomotor responsiveness [52, 53]. Fixation behavior was not different to controls in *upd2* knockdown flies: animals responded normally to two opposing ‘target’ bars, by walking decisively back and forth between them (measured by their low target deviation angle, see Methods) (Figure 6A). Optomotor behavior was also not significantly impacted in *upd2* knockdown animals (Figure 6B), suggesting these flies are able to perceive motion normally, along with being able to fixate on target objects. To investigate visual attention, we combined the two different kinds of visual stimuli (target and optomotor)in a visual competition scenario (Figure 6C), which allowed us to measure how much the moving grating distracted flies from the target stimulus [48, 49]. Surprisingly, in this attention paradigm *upd2* knockdown animals performed significantly better than controls, meaning that they were less distracted by the moving grating (Figure 6D).

**Fig 6.**
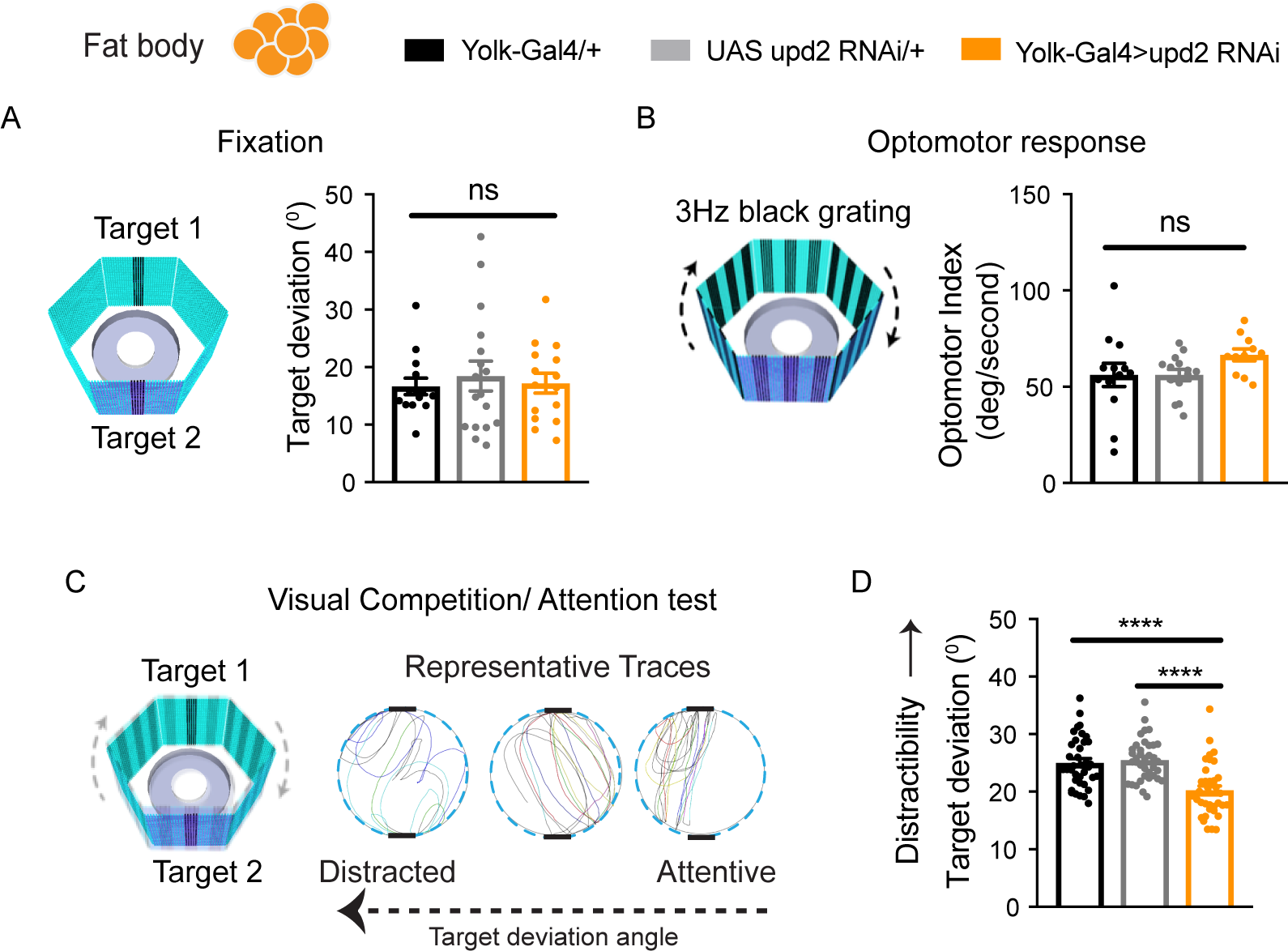
Visual attention behavior of upd2 deficient flies. (A) Fixation behavior in Buridan’s arena, where flies were presented with two opposing 7Hz flickering black bars. Fixation responses were calculated by target deviation in degrees and remained non-significant between genetic controls (grey and black bars) and knockdown flies (orange bar) (n=15) (B) Optomotor response to motion stimulus (3Hz black grating) determined by fly’s turning angle (^0^) per second for genetic controls (grey and black bars) and knockdown flies (orange bar) (n=11) (C) For visual competition (Attention) test flies were presented with a flickering bar (salient stimulus/target) and a 3Hz grating (distractor). The target deviation angle represents the measure of “distractibility”. Representative traces show the trajectories of single flies with low (left), average(middle) and high (right) target deviation angle. (D) Visual competition scores for knockdown flies (orange bar) were significantly lower compared to genetic controls, indicating less distractibility (n=21) One-way Anova with Tukey correction was used for comparing different conditions. *\*P <0.05, **P <0.01, ***P <0.001, ****P<0.0001 Error bars show SEM*.

*Upd2* mutants sleep less than control flies (Figures 1-4), suggesting that they are sleep deprived. However, sleep deprivation impairs attention in flies [27], which made us question whether the increased visual focus observed in *upd2*-deficient animals was a failure of attention, rather than improved attention. In other words, if attention is optimal in well-rested wild-type animals [54], a decreased capacity to detect a distracting stimulus (e.g., a moving grating) might be viewed as defective rather than improved attention. To address this conundrum, we decided to restore normal levels of sleep to *upd2* mutants, to see if this returned their visual attention to control levels. To increase sleep in *upd2* mutants, we used a sleep-promoting drug, THIP (4,5,6,7-tetrahydroisoxazolo-[5,4-c]pyridine-3-ol). Previous work has shown that THIP-induced sleep restores behavioral plasticity and attention to *Drosophila* mutants [16, 55], so we were curious if increased sleep in *upd2* would return distractibility to control levels, which would argue that the increased fixation observed in these animals was a *failure* of attention corrected by sufficient sleep. As expected, THIP exposure increased sleep in *upd2* mutants, up to similar levels as controls (Figure S6A). However, additional sleep in *upd2* knockdown animals did not change their visual attention phenotype compared to similarly-treated controls (Figure S6B). This suggests that the increased focus (or decreased distractibility) in these flies is a direct feature of their starved-like state rather than sub-optimal attention processes resulting from insufficient sleep.

We previously found that only *upd2* downregulation in the fat body decreased sleep, while downregulation in the muscle and nervous system had no effect on sleep (Figure 2). To determine whether this specificity extended to visual attention behavior as well, we next examined whether downregulation of *upd2* in other tissues (i.e., muscle and neurons) also altered visual attention. We found that *upd2* knockdown in muscle and neurons had no effect on visual behaviors or on visual attention (Figure S7A, B). Together, these results suggest that visual attention becomes more focused when the *Drosophila* fat body downregulates a satiety signal, and that this improved attention is preserved despite associated sleep loss.

### The *upd2* satiety signal regulates visual attention via the *domeless* receptor in insulin-expressing neurons

We next investigated how downregulation of *upd2* might be signaling a starvation signal to the fly brain. The *upd2* receptor, domeless (*dome*), is expressed in several subsets of neurons including mushroom body neurons, neuropeptide F (Npf) neurons and in PI region of the brain, where insulin-like peptide (Ilp) neurons are located [32, 56]. *Drosophila* Ilp neurons share functional similarities with mammalian insulin cells and are involved in nutrient sensing [57, 58]. Ilp-2, one of the three Ilps expressed in the insulin producing cells (IPC), (Figure 7A), have been previously shown to alter sleep and feeding [59, 60]. We therefore employed an RNAi strategy to downregulate dome in Ilp-2 neurons (Figure 7B). As before, we investigated sleep, feeding, and visual attention behaviors. Remarkably, these mirrored closely all of the *upd2* knockdown phenotypes: *dome* knockdown flies had a significant decrease in day-time sleep (Figure 7C-D); *dome* knockdown flies also displayed night-time hyperphagia (Figure 7E). However, their total food intake over 24 hours was not significantly different (Figure S8). Finally, when we investigated visual attention in *dome* knockdown animals, we saw significantly improved visual attention, compared to controls (Figure 7F), while there were no changes in their simple visual behaviors (Figure S9). These results suggest that the brain integrates the peripheral *upd2* signal secreted from the fat body via the *domeless* receptor on Ilp-2 neurons, to simultaneously decrease sleep need while sharpening visual attention.

**Fig 7.**
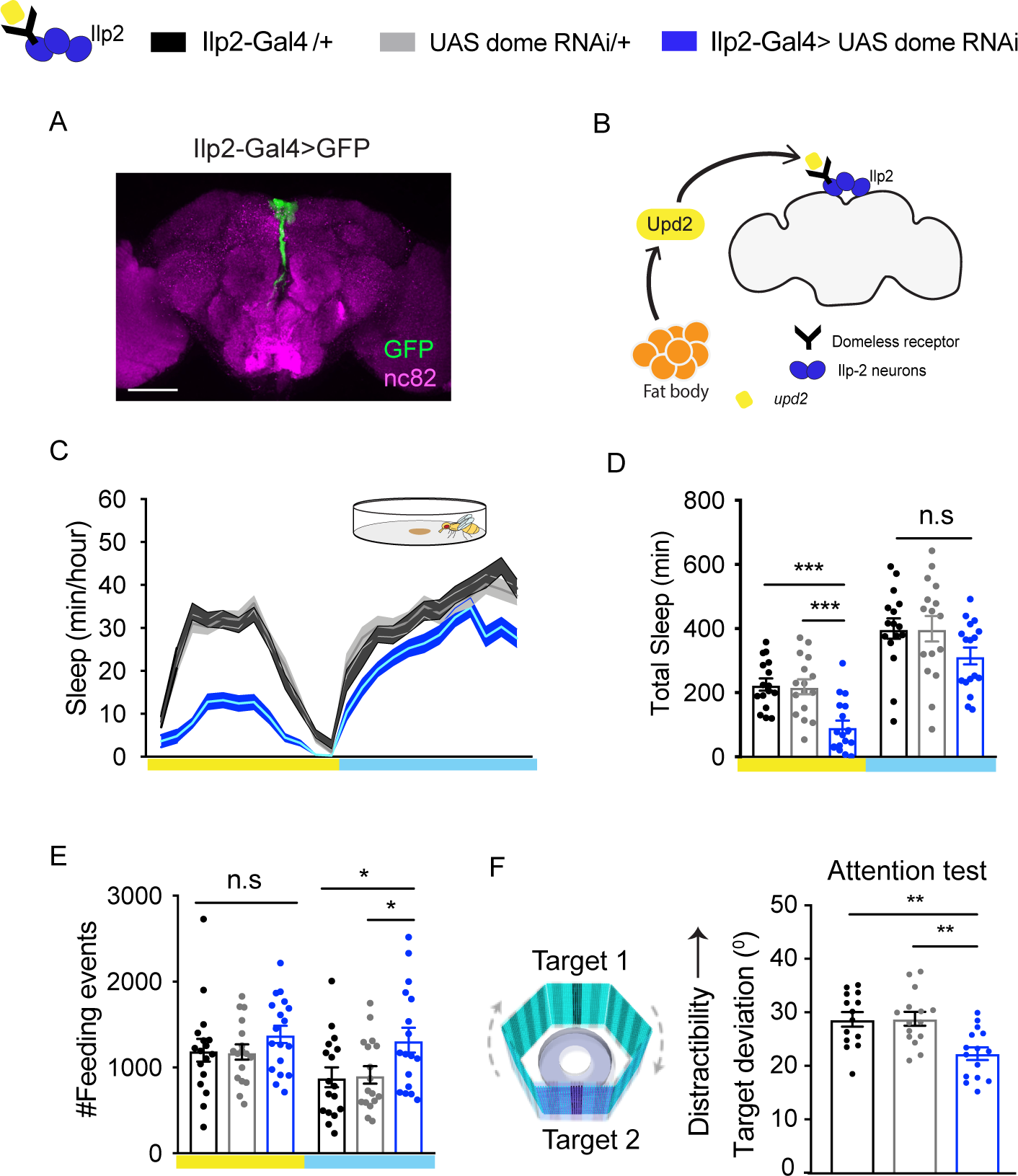
*Domeless,* the upd2 receptor in Insulin-like peptide 2 neurons integrates the peripheral satiety signal. (A) *Ilp2-GAL4* neurons are located in the pars intercerebralis (PI) region of the brain. Whole mount brain image, max projection, Scale bar 100 μm. (B) Schema illustrating the integration of *upd2* signal via domeless receptor in Ilp-2 neurons. (C) 24hr sleep profile of flies with knockdown of domeless receptor in Ilp2 neurons (blue) compared to parental controls (*Ilp2-GAL4/+*, black; *UAS domeRNAi/ +* (light grey). Sleep was measured in open-field arenas. (D) Day-time sleep was significantly reduced in knockdown flies, whereas night-time sleep remained was not affected. (E) Number of feeding events showed a significant increase at night-time. (F) Target deviation angle of flies with *dome* knockdown was significantly lower compared to genetic controls, indicating less distractibility (n=13-15). For sleep analysis 3-5day old adult females were used, n=15-17 per genotype. Sleep was recorded over 3 days. *\*P <0.05, **P <0.01, ***P <0.001, One-way ANOVA, Error bars show SEM*.

## Discussion

Sleep has been found to be necessary for maintaining cognitive properties such as memory, attention and decision making in several animals, including flies and humans [11,27,55,61–64]. This is because sleep most likely accomplishes a variety of important conserved functions, for any animal brain [41, 54]. It is therefore remarkable that sleep can under some circumstances be suppressed, as in the case of starvation. This suggests downstream pathways that can override some of the deleterious effects of sleep deprivation, especially effects relating to cognitive performance. That such mechanisms should exist seems adaptive: starving animals need to find food quickly, rather than sleep.

One important category of decisions animals make on a daily basis is their food choices, and how to source them. In humans, insufficient sleep has been found to alter desire for food and increase the preference for high caloric foods [65]. Additionally, nutrition strongly alters sleep amount and sleep quality [66]. Poor food choices are the most common cause of metabolic disorders such as type-2 diabetes and obesity [67, 68] and these disorders are also associated with sleep disorders [69, 70]. Yet, how sleep and feeding decisions influence each other is not well understood, and hard to disentangle in human patients. Manipulating satiety cues in animal models provides a way to establish some level of causality between sleep and feeding behavior.

A key roadblock toward addressing these questions in *Drosophila melanogaster* has been the lack of paradigms in which sleep quality and feeding behavior can be quantified in the same animals. Studying both behaviors simultaneously has been difficult given the small size of flies and their miniscule food intake. So far, the only platform which allowed this was the Activity Recording Capillary (ARC) Feeder [34, 71]. In our current study, we provide an alternative approach to measuring both behaviors by using a solid food source in open-field arenas amenable to visual tracking for sleep experiments. Our combined feeding and sleep paradigm confirmed a role for *upd2* in regulating feeding and sleep behavior in *Drosophila*. In future work, similar combined platforms (e.g., FlyPad [72]) could be employed to provide a more precise assessment of feeding behavior alongside sleep monitoring. This is crucial, as there is clearly substantial individual variability in feeding and sleep behavior (Figure 5B, Figure S4B).

Satiety requires signals from peripheral tissues to terminate food intake. The fat bodies are the main energy storage site in flies, and fulfills functions similar to mammalian adipose tissue and liver [35]. In *Drosophila*, *Upd2* is secreted from the fat bodies upon feeding [21]. Similarly, the mammalian ortholog leptin is secreted by adipose tissue, and is received as a satiety signal by the brain to inhibit appetite [22, 73]. Leptin levels in the blood change upon food intake [74] and are shown to fluctuate in a circadian manner [75]. Interestingly, a number of studies have found that leptin levels in rodents are significantly decreased in response to sleep deprivation [76–79]. On the other hand, leptin-deficient mice (Lep^ob/ob^) have been found to have altered sleep architecture, with increased non-rapid eye movement (NREM) sleep and shorter sleep bouts [80]. These studies therefore suggest a homeostatic relationship between leptin levels and sleep amount/quality. Our study shows that Upd2 in flies plays a similar role in sleep regulation, where lack of *upd2* reduces sleep duration and alters sleep quality. It is therefore probable that key sleep functions, for example that may be associated with deeper sleep, are not being satisfied in *upd2* mutants. Nevertheless, these animals have improved visual attention, and imposing more sleep on the mutants does not change their sharpened attention phenotype. This suggests that the satiety cue provided by *upd2* directly modulates attention-like behavior, regardless of sleep levels. As sleep deprivation normally impairs attention [27] and *upd2* mutants have reduced sleep, the effect of *upd2* on improving attention may ensure optimal cognitive performance in the face of suppressed sleep functions. In other words, if sleep is going to be sacrificed in order to promote foraging and feeding, then this behavior should be optimized rather than degraded. Although we did not examine foraging behavior in our study, our visual attention experiments probe a fundamental cognitive process (distractibility) that could affect many different kinds of behaviors, including foraging.

It is interesting to note that simple visual behaviors remained unaffected in genetically starved flies. Similarly, sleep deprivation does not seem to affect visual fixation and optomotor behavior in *Drosophila* [27]. Instead, the flies’ capacity to deal with competing visual stimuli is affected by sleep deprivation, as well as by a starvation cue, but in opposite directions. One interpretation of our results was that the decreased level of distractibility in *upd2* mutants reflects improved attention, which might be an adaptation towards finding appropriate food resources under sub-optimal nourishment conditions. While we have not completely excluded the possibility that this could be a form of impaired attention (to be less distractible can be maladaptive, as in autism), such a sharpened focus on innately attractive objects (for a walking fly, a dark bar is attractive [14]) could be seen as beneficial when animals are foraging for food.

Along with attention and sleep, another behavior that is significantly altered in *upd2* mutants is feeding itself, with night-time hyperphagia being a recurrent observation. Does decreased sleep quality alter feeding (and thus *upd2* levels), or do altered feeding patterns affect sleep quality? It is again difficult to disambiguate causality in this regard in humans, and even in animal models. In our *Drosophila* experiments we found an increase in R4 neuronal activity in *upd2* mutants, which correctly reflects their hunger status, as these neurons have been shown to respond under starvation regimes [43]. This suggests that a persistent hunger signal, reflected in R4 neuronal activity, may underlie the decreased sleep phenotype. This is further supported by our observation that prolonged activation of these neurons could decrease sleep. Considering the similarities between leptin/upd2 regulation, future studies temporally manipulating this hunger signal should be able disambiguate causal links between sleep need and nutritional status.

Our findings suggest that IIp2 cells are a key portal in the fly brain for integrating hunger signals and translating these into appropriate behavioral programs. How signaling from IIp2 cells in the fly brain leads to improved visual attention remains unknown. In fly larvae, Ilp2 is released within the brain and acts on a subset of neurons via insulin receptors [81]. So it is possible that, in adults, IIp2 cells target the ellipsoid body indirectly via insulin signaling, and that this would regulate selective attention by tuning circuits in the central complex that affect visual behavior more broadly [82, 83]. Alternatively, IIp2 cells might be communicating directly to arousal-regulatory circuits in the central complex, by way of gap junctions for example [84], to directly modulate behavioral responsiveness levels. Future experiments testing either of those possibilities should reveal the downstream mechanisms involved.

## Materials and Methods

### Fly stocks and maintenance

Flies were raised on standard agar-yeast based food at 25°C, 50-60% humidity with 12h light: 12h dark cycle. Starvation experiments were performed on 1% Agar food for 24 hours.

*upd2^Δ3-62^* was kindly provided by J. Hombria [30]. The UAS-CsChrimson strain was a gift from Vivek Jayaraman, Janelia Farm Research Campus, United States. Following stocks were from Bloomington stock center, yw (#1495), yolk-Gal4 (#58814), upd2-RNAi (#33988, HMS00901) [85], 24B-Gal4 (#1757), R57C10-Gal4 (#39171), R38H02-Gal4 (#47352) [86], UAS-CaLexa (#66542) [42]. Flies were crossed into a w+ background all experiments.

### Sleep and sleep intensity measurements

*Drosophila* arousal tracking (DART) software was used for sleep tracking and analysis [33]. 3-5-day old virgin females were placed into 65mm glass tubes (Trikinetics^TM^, Waltham, MA). A Logitech webcam (c9000 or c920) was used for sleep recordings with 5 frames per second. Sleep parameters were calculated according to traditional fly sleep criteria (Shaw et al. 2000; Hendricks et al. 2000), where sleep is defined as inactive durations for 5 minutes or more (a “sleep bout”). These were binned for every hour (“Sleep minutes/hour”). For sleep intensity measurements a vibration stimulus using motors (Precision Microdrives, 312-101) was presented every hour to measure behavioral responsiveness. Details of the software and calculations are described in Faville et. al., 2015.

For optogenetic experiments, flies were grown on 0.2 μM of all-trans-retinal (ATR) during the experiment and at least 1 day prior. Flies were exposed to low intensity white light from 8 am to 8 pm throughout the experiment, with additional red light illumination during Chrimson activation using Red-Orange LEDs (Luxeon Rebel, 617 nm, 700 mA, Phillips LXM2-PH021-0070) as previously [87]. Flies were illuminated with 9.5 µW/mm^2^ from four light-emitting diode (LED) arrays.

### Open-field behavioral analysis

The platform was custom made with white acrylic sheet. Each platform consisted of 36 individual chambers, with 36mm in diameter and 2.5 mm height. Center of the chamber had a hole with 3mm depth and 5mm width, where food was placed. The platform cover was covered with a transparent acrylic sheet, with 1mm holes surrounding each chamber.

Each food area was filled with a layer of 1% agar to increase the moisture of the food. Once solidified, it was covered with a layer of regular fly food.

Flies were transferred without anesthesia by using a mouth aspirator and acclimatized for minimum 12 hours prior to experiment. Fly sleep tracking and analyses were performed using DART [33] with custom-made MATLAB (Mathworks) scripts. Kinematic calculations were performed as previously described [88]. For feeding analysis the food area was detected using the Matlab function “imfindcircles” (object polarity was set to Dark). A fly was considered as feeding if it fulfilled 3 criteria: (1) distance from the feeding region (0mm from food pit), (2) speed= <1 mm/sec and (3) time spent feeding>30 seconds. The number of feeding events represent the total number of events throughout the recording.

### CApillary FEeding assay (CAFÉ)

The assay was slightly modified from the previously published versions (Ja *et al.*, 2007). 6-8 day old virgin female flies were used. Every testing chamber had 1% agar on the bottom, to eliminate the possibility of desiccation. Food (5% w/v sucrose, Sigma Aldrich) was presented in 5μL micropipettes (VWR, Westchester, PA) and the level of the meniscus was measured over time. For each condition 6-8 chambers were set, with 4-5 flies in each chamber. The experiments for different conditions were performed on the same day starting at ZT0-1, in an incubator with 25^0^C and 50-55% relative humidity. At least 5 empty chambers, without flies were used to control for the effects of evaporation.

### Buridan’s paradigm

A modified version of Buridan’s paradigm was used to assay for visual attention [48, 50]. Visual cues were presented on light emitting diode (LED) panels. Each LED panel contained 1024 individual LED units (32 rows by 32 columns) and was controlled via a LED Studio software (Shenzen Sinorad, Medical Electronics, Shenzen, China). Visual stimuli were created in Vision Egg software [89], written in Python programming language (L. Kirszenblat and Y. Zhou).

Three different visual cues were tested.

1. A moving grating (3 Hz) for testing the optomotor response behavior.
2. Two opposing flickering bars (7 Hz) for testing fixation behavior.
3. Competition stimulus (Figure-ground), with both grating and opposing flickering bars for testing selective visual attention.

Fixation and optomotor experiments lasted 1 minute. Figure-ground experiments lasted 3 minutes, during which the grating (clockwise or anticlockwise) was switched after 1.5 minutes.

For each test, female flies were collected as virgins and kept in vials in groups of 15-20 per vial. On day 2 their wings were cut under C0_2_ anesthesia and they were placed into fresh vials. They were given 2 days to recover from the effects of C0_2_ anesthesia. Tests were performed on a round platform (R=86mm), surrounded by a water-filled moat to prevent escape. The visual stimuli were alternated between each experiment from being presented on the horizontal or the vertical axes. Optomotor experiments were alternated between clockwise and anticlockwise gratings (1.5 minutes each). LED panels formed a hexagon, surrounding the platform (29cm diameter, 16cm height). The dark bar was 9 degrees in width and 45 degrees in height from the center of the arena. A camera (SONY Hi Resolution Colour Video Camera CCD-IRIS SSC-374) placed above the arena was used for tracking the movement of the fly on the platform at 30 frames per second. The open-source tracking software was used to record the position of the fly (Colomb et al. 2012)

Visual responses were analyzed by using CeTran (3.4) software (Colomb et al. 2012), and custom made scripts in R programming language (L.Kirszenblat and Y. Zhou). Target deviation was calculated as the smallest angle between the fly’s trajectory and the either of the vertical stripes. (Colomb et al. 2012). Optomotor index, was calculated as the angular velocity (turning angle/second) in the direction of the moving grating.

### Pharmacology

THIP (Gabaoxadol) was dissolved in standard food at 0.1 mg/ml for two days. For sleep experiments flies were transferred to tubes containing regular food. For visual behavior experiments, flies were transferred to regular food 1 hour prior to testing, as described previously[55].

### Immunohistochemistry and Confocal Imaging

Flies were collected under C0_2_ anesthesia and transferred to a drop of 1xPBS for dissection. After dissection, brains were transferred to a mini PCR-tube with 200 μl of 1xPBS. All of the following steps were performed on a rotator with 27rpm at room temperature. Brains were fixed with 4% paraformaldehyde diluted in PBS-T (1x PBS, 0.2 Triton-X 100) for 20-30min, followed by 3 washes in PBS-T. They were then blocked with 10%Goat serum (Sigma Aldrich, St. Louis, MO, USA) for 1h, followed by overnight primary antibody incubation. On the second day primary antibody was removed and brains were washed 3x with PBS-T. Then the secondary antibody was added and the tube was covered with aluminum foil for overnight incubation. On day three secondary antibody was removed and brains were washed with PBS-T. Primary antibodies were rabbit anti-GFP 1:1000 (Invitrogen), mouse anti-nc82 1:10 (DSHB). Secondary antibodies were anti-rabbit 488, 1:250 (Invitrogen), anti-mouse 647, 1:250 (Invitrogen). Brains were then transferred to microscope slides and mounted on a drop of Vectashield (Vector Laboratories, Burlingame, CA) for imaging. Images were acquired on a spinning-disk confocal system (Marianas; 3I, Inc.) consisting of an Axio Observer Z1 (Carl Zeiss) equipped with a CSU-W1 spinning-disk head (Yokogawa Corporation of America), ORCA-Flash4.0 v2 sCMOS camera (Hamamatsu Photonics), 20x 0.8 NA PlanApo and 100x 1.4 NA PlanApo objectives were used and image acquisition was performed using SlideBook 6.0 (3I, Inc).

For CaLexA experiments same acquisition settings were used between different conditions. Fiji (ImageJ) was used for image processing. GFP intensity measurements were done using Fiji intensity measurement plug-in.

### Statistical Analyses

Statistical analyses were performed using Prism 7.0a (GraphPad Software, Inc). Normality tests were performed using Shapiro-Wilk normality tests. For normally distributed data two-tailed, unpaired Students t-test or one-way ANOVA, followed by Tukey correction was performed. Unless otherwise stated all data-sets represent mean + SEM.

## Acknowledgements

We would like to thank the Goetz lab for antibodies. Imaging was performed at the Queensland Brain Institute’s Advanced Microscopy Facility using Yokogawa spinning disk confocal. We thank Burczyk/Faville/Kottler (BFK) for adjustments made to the DART software for tracking feeding behavior.

Conceptualization: D.E, BvS

Data curation: D.E, L.K

Formal analysis: D.E

Funding acquisition: BvS

Investigation: D.E, L.K

Methodology: D.E, L.K

Project administration: D.E, BvS

Resources: BvS

Software: R.F Supervision: D.E, BvS

Validation: D.E

Visualization: D.E

Writing – original draft preparation: D.E, L.K, BvS

Writing – review & editing: D.E, L.K, BvS

## Supporting Information

**S1 Fig.**
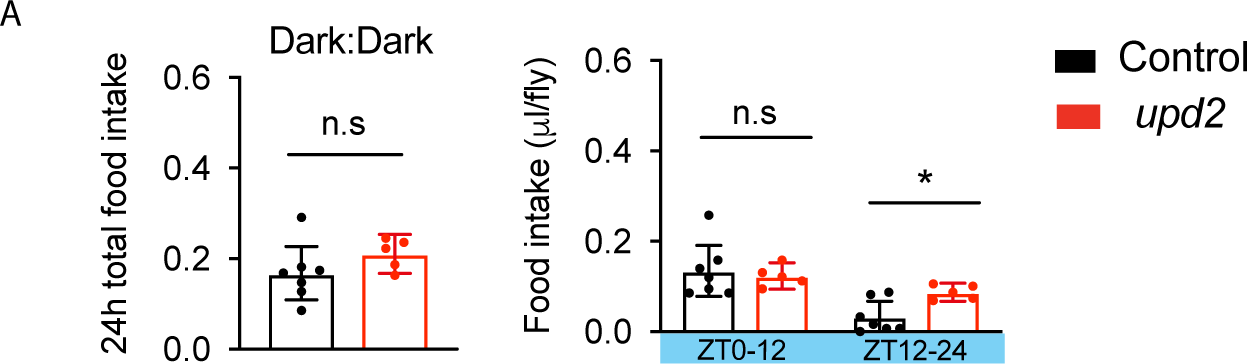
Night-time hyperphagia of *upd2* mutants is not dependent on light entrainment. (A) Scores for feeding experiments under constant darkness (DD). *Upd2* mutants (red) had decreased night time food intake compared to controls (black) (n=25-35 flies with 5 flies per Café chamber). *\*P <0.05, **P <0.01, ***P <0.001, Student’s t-test, Error bars show SEM*.

**S2 Fig.**
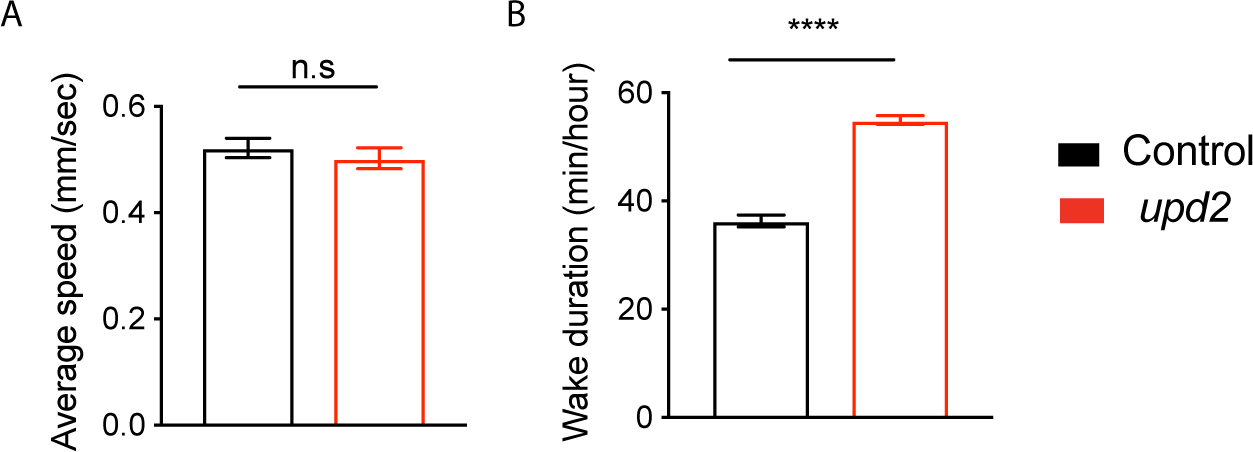
*Upd2* mutant flies are hyperactive yet have comparable speed to controls. (A) Average speed of *upd2* mutant flies (red) was not significantly different to controls (black). (B) Mutant flies had increased wake duration compared to controls. *\*P <0.05,**P <0.01, ***P <0.001,* Analyses in this figure is from the same dataset as in Figure 1. *Student’s t-test for normally distributed data or Mann-Whitney U rank-sum test for nonparametric data was used to compare datasets. *P <0.05, **P <0.01, ***P <0.001, Error bars show SEM*.

**S3 Fig.**
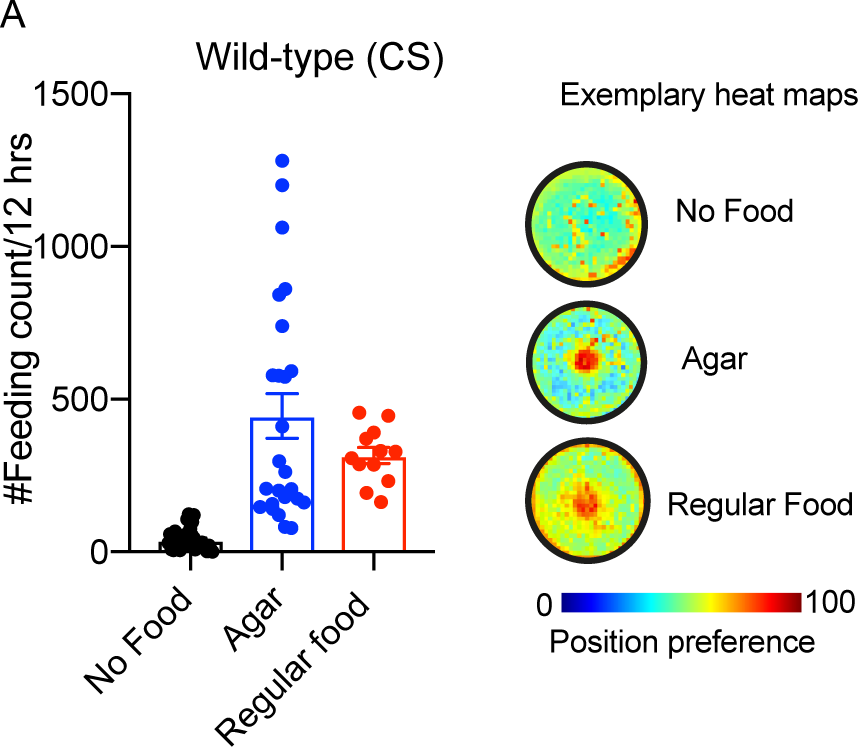
Control experiments for feeding tracking. (A) Under no food conditions, food area was covered with parafilm to prevent access to food (Black bar). Flies did not spend time in the center in the absence of food. When presented with agar only, they spent a similar time as they did with regular food (Blue bar) Exemplary heat maps for flies under different food conditions (Left panel).

**S4 Fig.**
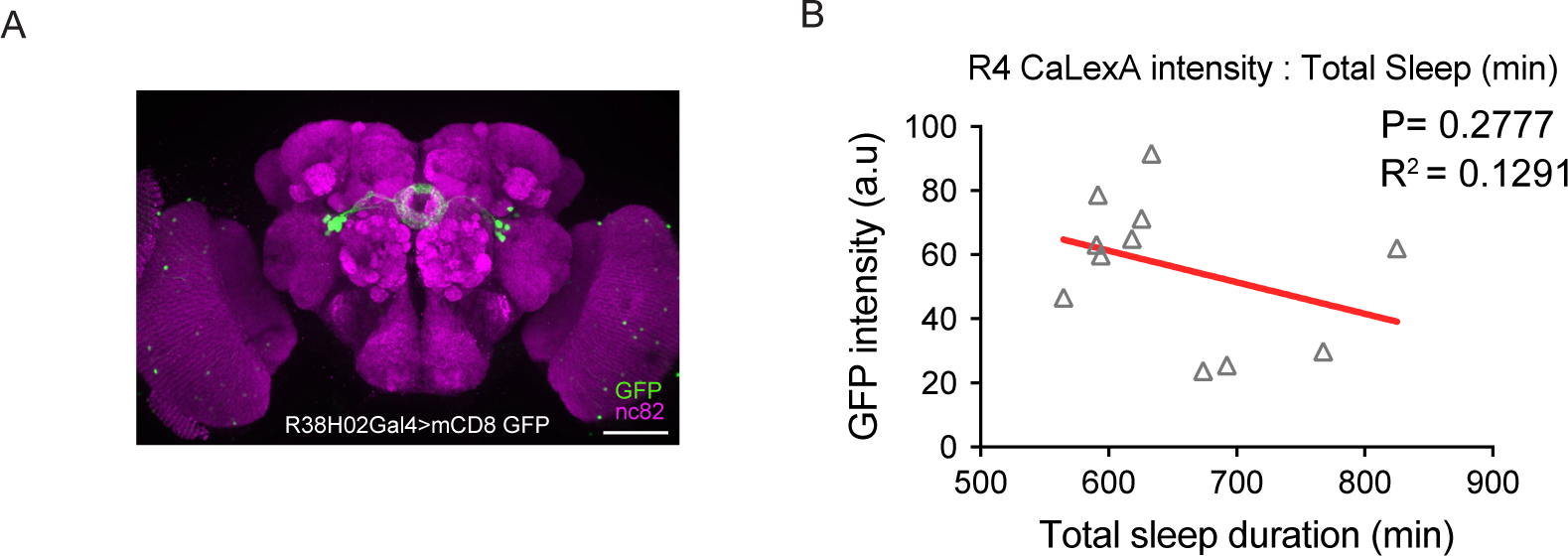
Baseline R4 neuron activity does not correlate with sleep duration. (A) Expression of R38H02-Gal4 in the brain using UAS-mCD8 GFP (green). Neuropil is stained with bruchpilot (magenta) (Scale bar: 100 µm) (B) CaLexA intensity of individual flies are plotted against their total sleep duration (24h). Flies were housed in open-field DART arena. CaLexA intensity did not correlate sleep duration. Two-tailed p-values for Pearson’s correlation coefficient are shown. Analyses in this figure is from the same dataset as in Figure 4.

**S5 Fig.**
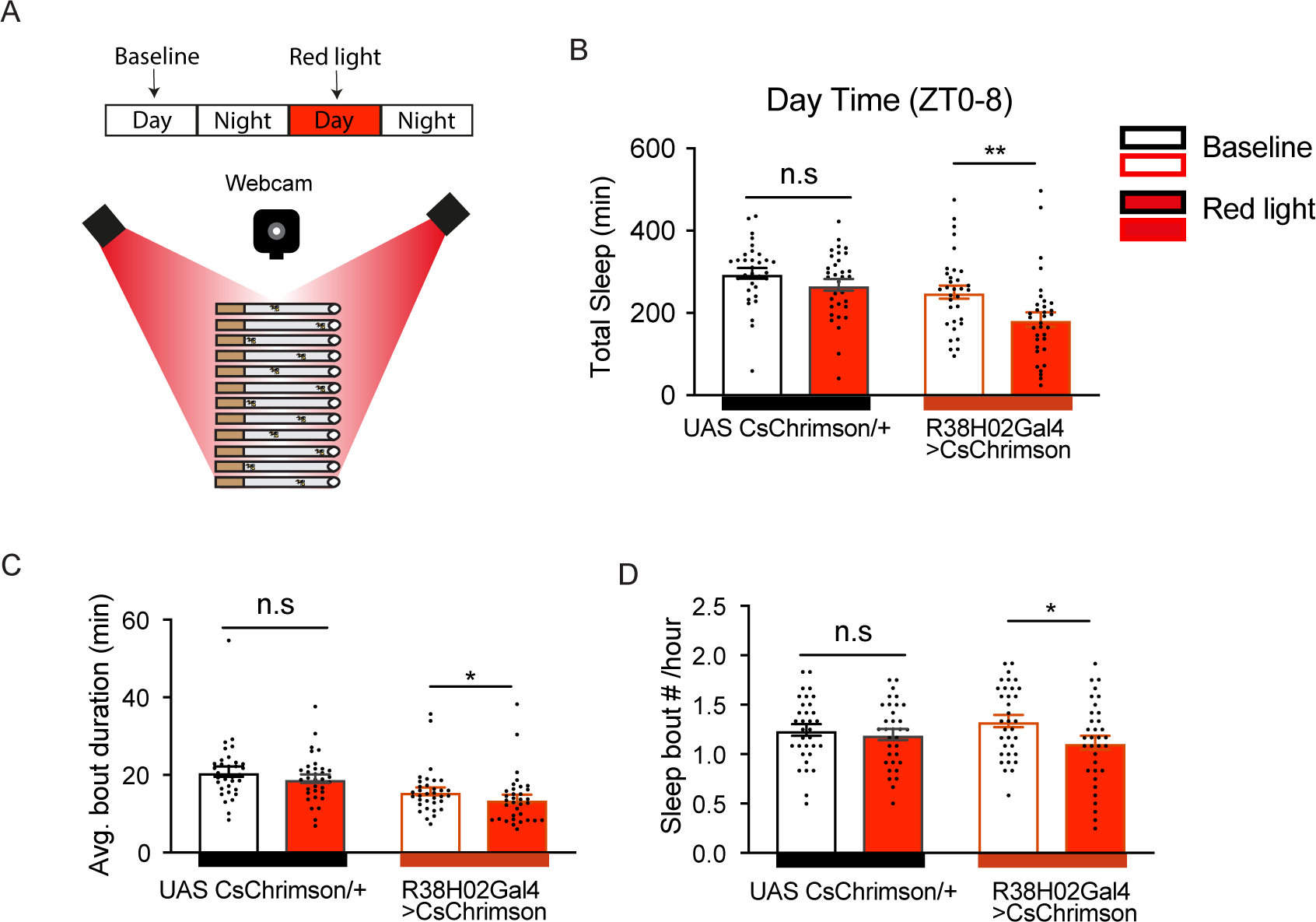
Optogenetic activation of R4 neurons decreases sleep. (A) Flies expressing a red-light-activated channelrhodopsin (Chrimson) were placed in to tubes and sleep was analyzed under baseline conditions on day 1 (normal white light from 8 am to 8 pm, followed by 12 hours darkness at night), an activated condition on day 2 (normal white light + 12 hours red light illumination from 8 am to 8 pm, followed by 12 hours darkness at night. (B) Compared to baseline day optogenetic activation of R4 neurons, via R38H02 Gal4, led to a significant decrease in sleep duration (right panel) but UAS CsChrimson/+ controls showed no difference between baseline and activation day (left panel). (C-D) Both bout duration and bout number were also significantly reduced under red light activation in R38H02-Gal4> CsChrimson flies. Red light activation had no effect on the bout duration or bout number of control flies (UAS CsChrimson/+, left panels). (n=34-35 flies), *One-way Anova (with Tukey’s post-hoc test) for normally distributed data or Kruskal-Wallis test (with Dunn’s multiple comparison test) for nonparametric data was used to compare datasets. *P <0.05, **P <0.01, ***P <0.001, Error bars show SEM*.

**S6 Fig.**
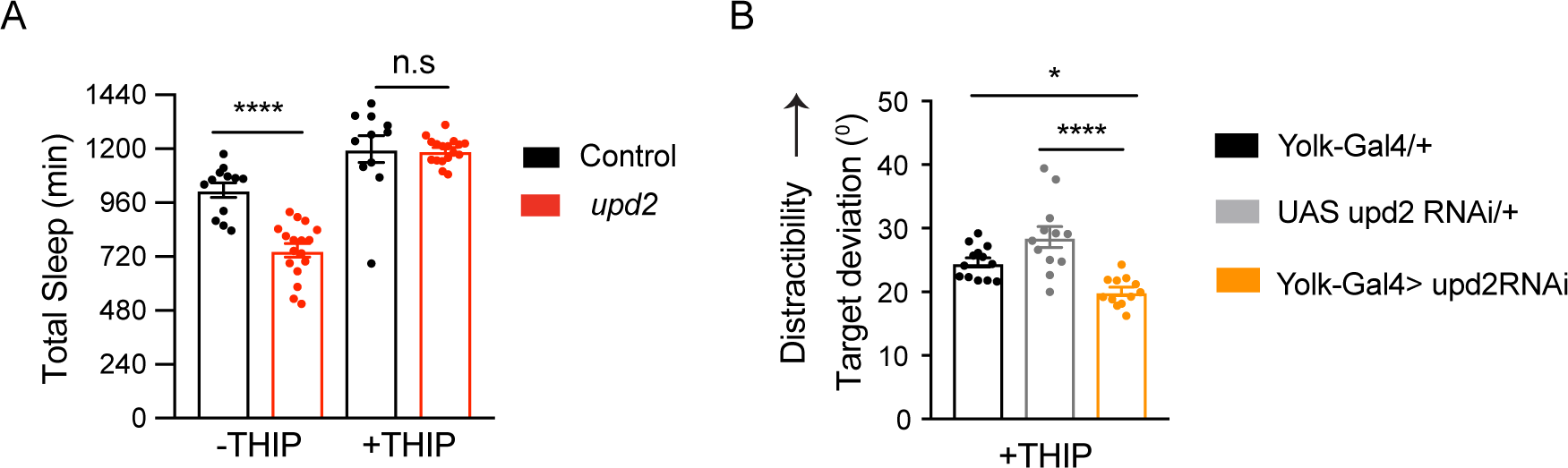
Gaboxadol induced sleep does not rescue attention phenotype. (A) Non-THIP fed *upd2* mutants (red,left) had significantly reduced sleep compared to controls (black). THIP (0.1 mg/ml) fed *upd2* mutants and control flies sleep sleep was not significantly different (n=12-17), *Student’s t-test, Error bars show SEM.* (B) Sleep induction via THIP did not rescue the increased attention phenotype of flies with fat body upd2 knockdown (orange, Yolk Gal4> UAS upd2 RNAi) compared to controls (Yolk-Gal4/+, black and UAS upd2 RNAi/+, grey. (n=12-13) One-way Anova with Tukey correction was used for comparing different conditions. *\*P <0.05, **P <0.01, ***P <0.001, ****P<0.0001 Error bars show SEM*.

**S7 Fig.**
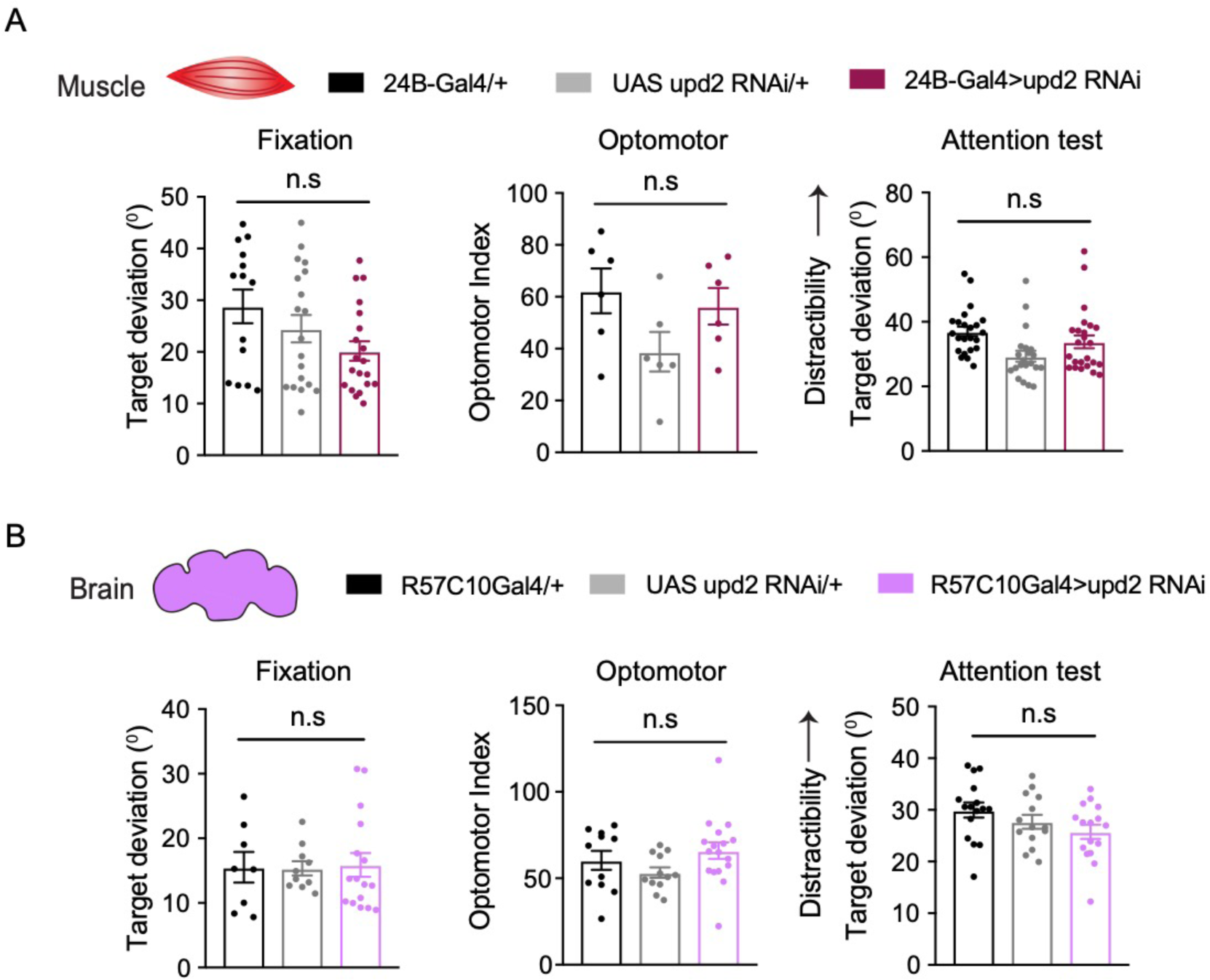
Knockdown of *upd2* in muscles or pan neuronally has no effect on visual attention. (A) We did not observe any differences in simple visual behaviors (Fixation (n=15-20), optomotor (n=6-8) or in visual attention (n=13-16) flies with muscle specific *upd2* knockdown (24B-Gal4> UAS upd2 RNAi, maroon) compared to controls (24B-Gal4/+, black and UAS upd2 RNAi/+, grey). (B) Pan-neuronal knockdown of *upd2* (R57C10-Gal4> UAS upd2 RNAi, purple) also had no impact on visual (optomotor, n=12-16), fixation n=8-16) and visual attention behaviors, (n= 14-16) compared to genetic controls (R57C10-Gal4/+, black and UAS upd2 RNAi, grey). One-way Anova with Tukey correction was used for comparing different conditions. *\*P <0.05, **P <0.01, ***P <0.001, ****P<0.0001 Error bars show SEM*.

**S8 Fig.**
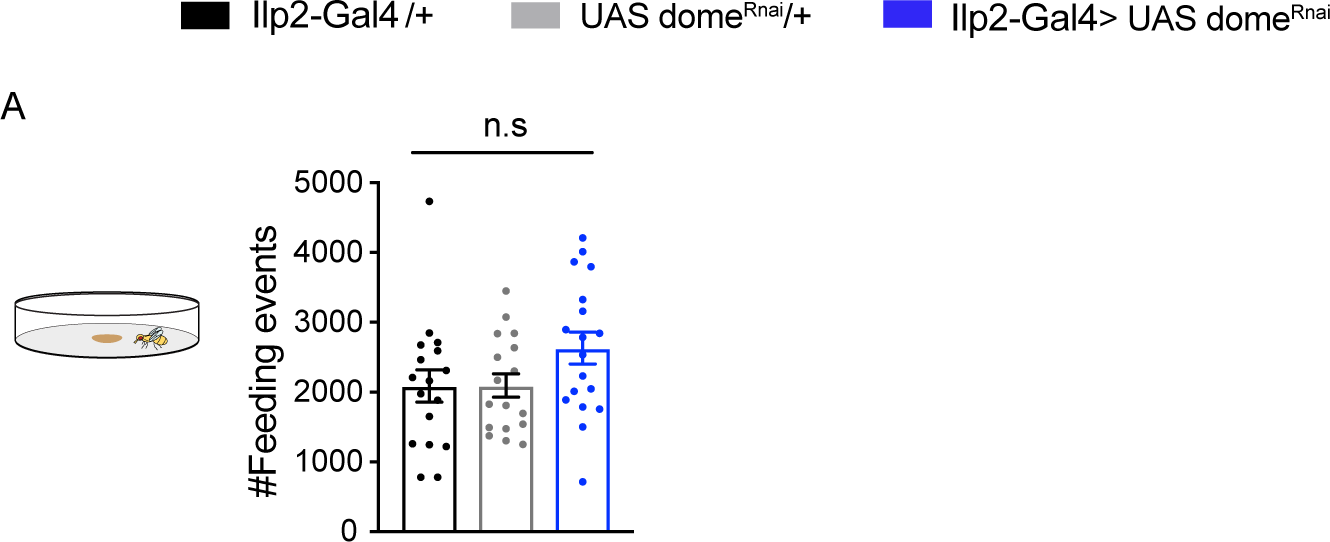
*Domeless* knockdown does not alter total food intake. (A) Number of total feeding events was not significantly different between control flies and domeless knockdown flies. Analyses in this figure is from the same dataset as in Figure 7D-E. One-way Anova with Tukey correction was used for comparing different conditions. *Error bars show SEM*.

**S9 Fig.**
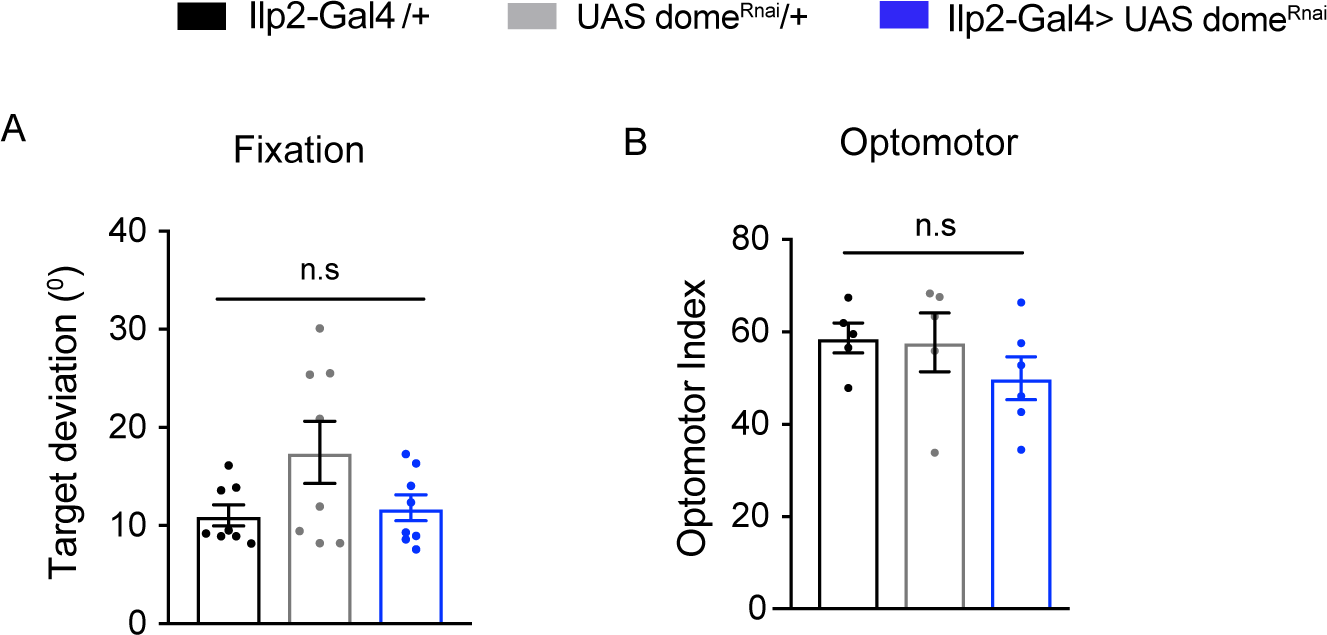
*Domeless* knockdown does not alter simple visual behaviors. (A) Fixation and (B) optomotor behavior of Ilp2-Gal4> UAS dome RNAi (blue) were not significantly different to Ilp2-Gal4/+, black and UAS dome RNAi/+, grey. n=6-8 per experiment, One-way Anova with Tukey correction was used for comparing different conditions. *Error bars show SEM*.

## References

1. Keene AC, Duboue ER. The origins and evolution of sleep. J Exp Biol. 2018;221: jeb159533. doi:10.1242/jeb.159533

2. Campbell SS, Tobler I. Animal sleep: A review of sleep duration across phylogeny. Neurosci Biobehav Rev. 1984;8: 269–300. doi:10.1016/0149-7634(84)90054-X

3. Siegel JM. Clues to the functions of mammalian sleep. Nature. 2005;437: 1264–1271. doi:10.1038/nature04285

4. Krause AJ, Simon E Ben, Mander BA, Greer SM, Saletin JM, Goldstein-Piekarski AN, et al. The sleep-deprived human brain. Nat Rev Neurosci. 2017;18: 404–418. doi:10.1038/nrn.2017.55

5. Andermann ML, Lowell BB. Toward a Wiring Diagram Understanding of Appetite Control. Neuron. 2017;95: 757–778. doi:10.1016/j.neuron.2017.06.014

6. Sternson SM, Atasoy D, Betley JN, Henry FE, Xu S. Review An Emerging Technology Framework for the Neurobiology of Appetite. Cell Metab. 2016;23: 234–253. doi:10.1016/j.cmet.2015.12.002

7. Sternson SM, Eiselt A-K. Three Pillars for the Neural Control of Appetite. Annu Rev Physiol. 2017;79: 401–423. doi:10.1146/annurev-physiol-021115-104948

8. Mugnai G, Danese A. Sleep Deprivation and Metabolic Syndrome [Internet]. Modulation of Sleep by Obesity, Diabetes, Age, and Diet. Elsevier Inc.; 2014. doi:10.1016/B978-0-12-420168-2.00020-X

9. Koren D, Dumin M, Gozal D. Diabetes, Metabolic Syndrome and Obesity: Targets and Therapy Dovepress Role of sleep quality in the metabolic syndrome. 2016; 281–310. doi:10.2147/DMSO.S95120

10. Pool AH, Scott K. Feeding regulation in Drosophila. Curr Opin Neurobiol. 2014;29: 57–63. doi:10.1016/j.conb.2014.05.008

11. Donlea JM. Roles for sleep in memory: insights from the fly. Curr Opin Neurobiol. 2019;54: 120–126. doi:10.1016/j.conb.2018.10.006

12. Hendricks JC, Finn SM, Panckeri KA, Chavkin J, Williams JA, Sehgal A, et al. Rest in Drosophila Is a Sleep-like State. Neuron. 2000;25: 129–138. doi:10.1016/S0896-6273(00)80877-6

13. Shaw PJ, Cirelli C, Greenspan RJ, Tononi G. Correlates of sleep and waking in Drosophila melanogaster. Science. 2000;287: 1834–1837. doi:10.1126/science.287.5459.1834

14. Grabowska MJ, Steeves J, Alpay J, Van De Poll M, Ertekin D, van Swinderen B. Innate visual preferences and behavioral flexibility in Drosophila. J Exp Biol. 2018;221: jeb185918. doi:10.1242/jeb.185918

15. Paulk AC, Kirszenblat L, Zhou Y, van Swinderen B. Closed-Loop Behavioral Control Increases Coherence in the Fly Brain. J Neurosci. 2015;35: 10304–15. doi:10.1523/JNEUROSCI.0691-15.2015

16. Kirszenblat L, Yaun R, van Swinderen B. Visual experience drives sleep need in Drosophila. Sleep. 2019; 1–12. doi:10.1093/sleep/zsz102

17. Keene AC, Duboué ER, McDonald DM, Dus M, Suh GSB, Waddell S, et al. Clock and cycle limit starvation-induced sleep loss in drosophila. Curr Biol. 2010;20: 1209–1215. doi:10.1016/j.cub.2010.05.029

18. Catterson JH, Knowles-Barley S, James K, Heck MMSS, Harmar AJ, Hartley PS. Dietary modulation of Drosophila sleep-wake behaviour. PLoS One. 2010;5: e12062. doi:10.1371/journal.pone.0012062

19. Masek P, Reynolds L a, Bollinger WL, Moody C, Mehta A, Murakami K, et al. Altered regulation of sleep and feeding contribute to starvation resistance in Drosophila. J Exp Biol. 2014;217: 3122–3132. doi:10.1242/jeb.103309

20. Lee G, Park JH. Hemolymph sugar homeostasis and starvation-induced hyperactivity affected by genetic manipulations of the adipokinetic hormone-encoding gene in Drosophila melanogaster. Genetics. 2004;167: 311–323. doi:10.1534/genetics.167.1.311

21. Rajan A, Perrimon N. Drosophila cytokine unpaired 2 regulates physiological homeostasis by remotely controlling insulin secretion. Cell. 2012;151: 123– 137. doi:10.1016/j.cell.2012.08.019

22. Londraville RL, Prokop JW, Duff RJ, Liu Q, Tuttle M. On the molecular evolution of leptin, leptin receptor, and endospanin. Front Endocrinol (Lausanne). 2017;8. doi:10.3389/fendo.2017.00058

23. Frederich RC, Hamann A, Anderson S, Lollmann B, Lowell BB, Flier JS. Leptin levels reflect body lipid content in mice: evidence for diet-induced resistance to leptin action. Nat Med. 1995;1: 1311–1314.

24. Liu S, Liu Q, Tabuchi M, Wu MN. Sleep drive is encoded by neural plastic changes in a dedicated circuit. Cell. 2016;165: 1347–1360. doi:10.1016/j.cell.2016.04.013

25. Seugnet L, Suzuki Y, Vine L, Gottschalk L, Shaw PJ. D1 Receptor Activation in the Mushroom Bodies Rescues Sleep-Loss-Induced Learning Impairments in Drosophila. Curr Biol. 2008;18: 1110–1117. doi:10.1016/j.cub.2008.07.028

26. Ganguly-Fitzgerald I, Donlea J, Shaw PJ. Waking Experience Affects Sleep Need in Drosophila. Science. 2006;313: 1775–1781. doi:10.1126/science.1130408

27. Kirszenblat L, Ertekin D, Goodsell J, Zhou Y, Shaw PJ, van Swinderen B. Sleep regulates visual selective attention in Drosophila. J Exp Biol. 2018;221: jeb191429. doi:10.1242/jeb.191429

28. Donlea J, Leahy A, Thimgan MS, Suzuki Y, Hughson BN, Sokolowski MB, et al. foraging alters resilience/vulnerability to sleep disruption and starvation in Drosophila. Proc Natl Acad Sci. 2012;109: 2613–2618. doi:10.1073/pnas.1112623109

29. Thimgan MS, Suzuki Y, Seugnet L, Gottschalk L, Shaw PJ. The Perilipin homologue, Lipid storage droplet 2, regulates sleep homeostasis and prevents learning impairments following sleep loss. PLoS Biol. 2010;8: 29–30. doi:10.1371/journal.pbio.1000466

30. Hombría JC-GG, Brown S, Häder S, Zeidler MP. Characterisation of Upd2, a Drosophila JAK/STAT pathway ligand. Dev Biol. 2005;288: 420–433. doi:10.1016/j.ydbio.2005.09.040

31. Ja WW, Carvalho GB, Mak EM, de la Rosa NN, Fang AY, Liong JC, et al. Prandiology of Drosophila and the CAFE assay. Proc Natl Acad Sci U S A. 2007;104: 8253–8256. doi:0702726104 [pii]\r10.1073/pnas.0702726104

32. Beshel J, Dubnau J, Beshel J, Dubnau J, Zhong Y. A Leptin Analog Locally Produced in the Brain Acts via a Conserved Neural Circuit to Modulate. Cell Metab. 2017;25: 208–217. doi:10.1016/j.cmet.2016.12.013

33. Faville R, Kottler B, Goodhill GJ, Shaw PJ, Swinderen B Van, van Swinderen B. How deeply does your mutant sleep? Probing arousal to better understand sleep defects in Drosophila. Sci Rep. 2015;5: 8454. doi:10.1038/srep08454

34. Murphy KR, Deshpande SA, Yurgel ME, Quinn JP, Weissbach JL, Keene AC, et al. Postprandial sleep mechanics in Drosophila. Elife. 2016; 1–19. doi:10.7554/eLife.19334

35. Arrese EL, Soulages JL. Insect fat body: energy, metabolism, and regulation. Annu Rev Entomol. 2010;55: 207–225. doi:10.1146/annurev-ento-112408-085356

36. Canavoso LE, Jouni ZE, Karnas KJ, Pennington JE, Wells MA. Fat metabolism in insects. Annu Rev Nutr. 2001;21: 23–46. doi:10.1146/annurev.nutr.21.1.23

37. Georgel P, Naitza S, Kappler C, Ferrandon D, Zachary D, Swimmer C, et al. Drosophila Immune Deficiency (IMD) Is a Death Domain Protein that Activates Antibacterial Defense and Can Promote Apoptosis. Dev Cell. 2001;1: 503– 514. doi:10.1016/S1534-5807(01)00059-4

38. Zhao X, Karpac J. Muscle Directs Diurnal Energy Homeostasis through a Myokine-Dependent Hormone Module in Article Muscle Directs Diurnal Energy Homeostasis through a Myokine-Dependent Hormone Module in Drosophila. Curr Biol. 2017;27: 1941–1955.e6. doi:10.1016/j.cub.2017.06.004

39. Brand AH, Perrimon N. Targeted gene expression as a means of altering cell fates and generating dominant phenotypes. Development. 1993;118: 401–415.

40. Jenett A, Rubin GM, Ngo T-TTB, Shepherd D, Murphy C, Dionne H, et al. A GAL4-Driver Line Resource for Drosophila Neurobiology. Cell Rep. 2012;2: 991–1001. doi:10.1016/j.celrep.2012.09.011

41. Kirszenblat L, van Swinderen B. Chapter 22 - Sleep in Drosophila. Handbook of Sleep Research. Elsevier B.V.; 2019. pp. 333–347. doi:https://doi.org/10.1016/B978-0-12-813743-7.00022-0

42. Masuyama K, Zhang Y, Rao Y, Wang JW. Mapping Neural Circuits with Activity-Dependent Nuclear Import of a Transcription Factor. J Neurogenet. 2012;26: 89–102. doi:10.3109/01677063.2011.642910

43. Dus M, Ai M, Suh GSB. Taste-independent nutrient selection is mediated by a brain-specific Na+ /solute co-transporter in Drosophila. Nat Neurosci. 2013;16: 526–8. doi:10.1038/nn.3372

44. Seugnet L, Suzuki Y, Thimgan M, Donlea J, Gimbel SI, Gottschalk L, et al. Identifying sleep regulatory genes using a Drosophila model of insomnia. J Neurosci. 2009;29: 7148–57. doi:10.1523/JNEUROSCI.5629-08.2009

45. Krashes MJ, DasGupta S, Vreede A, White B, Armstrong JD, Waddell S. A Neural Circuit Mechanism Integrating Motivational State with Memory Expression in Drosophila. Cell. 2009;139: 416–427. doi:10.1016/j.cell.2009.08.035

46. Ledue EE, Mann K, Koch E, Chu B, Dakin R, Gordon MD, et al. Starvation-Induced Depotentiation of Bitter Taste in Starvation-Induced Depotentiation of Bitter Taste in Drosophila. Curr Biol. 2016; 1–8. doi:10.1016/j.cub.2016.08.028

47. Sayin S, De Backer J-F, Siju KP, Wosniack ME, Lewis LP, Frisch L-M, et al. A Neural Circuit Arbitrates between Persistence and Withdrawal in Hungry Drosophila. Neuron. 2019; 1–15. doi:10.1016/j.neuron.2019.07.028

48. Colomb J, Reiter L, Blaszkiewicz J, Wessnitzer J, Brembs B. Open source tracking and analysis of adult Drosophila locomotion in Buridan’s paradigm with and without visual targets. PLoS One. 2012;7: 1–12. doi:10.1371/journal.pone.0042247

49. Ferguson L, Petty A, Rohrscheib C, Troup M, Kirszenblat L, Eyles DW, et al. Transient Dysregulation of Dopamine Signaling in a Developing Drosophila Arousal Circuit Permanently Impairs Behavioral Responsiveness in Adults. Front Psychiatry. 2017;8: 22. doi:10.3389/fpsyt.2017.00022

50. Götz KG. Visual Guidance in Drosophila. In: Siddiqi O, Babu P, Hall LM, Hall JC, editors. Development and Neurobiology of Drosophila. Boston, MA: Springer US; 1980. pp. 391–407. doi:10.1007/978-1-4684-7968-3_28

51. . Heisenberg, Martin and Wolf R. Vision in Drosophila: genetics of microbehavior. Springer-Verlag; 1984.

52. Gotz KG, Wandel U. Optomotor control of the force of flight in Drosophila and Musca. Biol Cybern. 1984;51: 135–139. doi:10.1007/bf00357927

53. Gotz KG. Optomoter studies of the visual system of several eye mutants of the fruit fly Drosophila. Kybernetik. 1964;2: 77–92.

54. Kirszenblat L, van Swinderen B. The Yin and Yang of Sleep and Attention. Trends Neurosci. 2015;38: 776–786. doi:10.1016/j.tins.2015.10.001

55. Dissel S, Angadi V, Kirszenblat L, Suzuki Y, Donlea J, Klose M, et al. Sleep restores behavioral plasticity to drosophila mutants. Curr Biol. 2015;25: 1270–1281. doi:10.1016/j.cub.2015.03.027

56. Brown S, Hu N, Hombria JC. Identification of the first invertebrate interleukin JAK/STAT receptor, the Drosophila gene domeless. Curr Biol. 2001;11: 1700– 1705.

57. Rulifson EJ, Kim SK, Nusse R. Ablation of insulin-producing neurons in flies: growth and diabetic phenotypes. Science. 2002;296: 1118–1120. doi:10.1126/science.1070058

58. Broughton S, Partridge L. Insulin/IGF-like signalling, the central nervous system and aging. Biochem J. 2009;418: 1–12. doi:10.1042/bj20082102

59. Cong X, Wang H, Liu Z, He C, An C, Zhao Z. Regulation of Sleep by Insulin-like Peptide System in Drosophila melanogaster. Sleep. 2015;38. doi:10.5665/sleep.4816

60. Metaxakis A, Tain LS, Grönke S, Hendrich O, Hinze Y, Birras U, et al. Lowered Insulin Signalling Ameliorates Age-Related Sleep Fragmentation in Drosophila. PLoS Biol. 2014;12. doi:10.1371/journal.pbio.1001824

61. Seugnet L, Dissel S, Thimgan M, Cao L, Shaw PJ. Identification of genes that maintain behavioral and structural plasticity during sleep loss. Front Neural Circuits. 2017;11: 1–16. doi:10.3389/fncir.2017.00079

62. Landolt H-P, Sousek A, Camillo HS. Effects of acute and chronic sleep deprivation. ESRS Eur Sleep Med Textb. 2014; 49–62.

63. Moraes DA, Venancio DP, Suchecki D. Sleep deprivation alters energy homeostasis through non-compensatory alterations in hypothalamic insulin receptors in Wistar rats. Horm Behav. 2014;66: 705–712. doi:10.1016/j.yhbeh.2014.08.015

64. Van Cauter E, Spiegel K, Tasali E, Leproult R. Metabolic consequences of sleep and sleep loss. Sleep Med. 2008;9 Suppl 1: S23–8. doi:S1389-9457(08)70013-3 [pii]\r10.1016/S1389-9457(08)70013-3

65. Greer SM, Goldstein AN, Walker MP. The impact of sleep deprivation on food desire in the human brain. Nat Commun. 2013;4: 2259. doi:10.1038/ncomms3259

66. Theorell-haglöw J, Lindberg E. Sleep Duration and Obesity in Adults: What Are the Connections ? Curr Obes Rep. 2016; 333–343. doi:10.1007/s13679- 016-0225-8

67. Hu FB, Manson JE, Stampfer MJ, Colditz G, Liu S, Solomon CG, et al. Diet, lifestyle, and the risk of type 2 diabetes mellitus in women. N Engl J Med. 2001;345: 790–797. doi:10.1056/NEJMoa010492

68. Schwingshackl L, Schwedhelm C, Hoffmann G, Knüppel S, Iqbal K, Andriolo V, et al. Food Groups and Risk of Hypertension: A Systematic Review and Dose-Response Meta-Analysis of Prospective Studies. Adv Nutr An Int Rev J. 2017;8: 793–803. doi:10.3945/an.117.017178

69. Grandner MA, Seixas A, Shetty S, Shenoy S. Sleep Duration and Diabetes Risk: Population Trends and Potential Mechanisms. Curr Diab Rep. 2016; doi:10.1007/s11892-016-0805-8

70. Nedeltcheva A V., Scheer FAJL. Metabolic effects of sleep disruption, links to obesity and diabetes. Curr Opin Endocrinol Diabetes Obes. 2014;21: 293–298. doi:10.1097/MED.0000000000000082

71. Murphy KR, Park JH, Huber R, Ja WW. Simultaneous measurement of sleep and feeding in individual Drosophila. Nat Protoc. 2017;12: 2355–2366. doi:10.1038/nprot.2017.096

72. Itskov PM, Moreira J-M, Vinnik E, Lopes G, Safarik S, Dickinson MH, et al. Automated monitoring and quantitative analysis of feeding behaviour in Drosophila. Nat Commun. 2014;5: 4560. doi:10.1038/ncomms5560

73. Elias CF, Aschkenasi C, Lee C, Kelly J, Ahima RS, Bjorbæk C, et al. Leptin differentially regulates NPY and POMC neurons projecting to the lateral hypothalamic area. Neuron. 1999;23: 775–786. doi:10.1016/S0896- 6273(01)80035-0

74. Shea SA, Hilton MF, Orlova C, Ayers RT, Christos S. Independent Circadian and Sleep/Wake Regulation of Adipokines and Glucose in Humans Steven. J Clin Endocrinol Metab. 2005;90: 2537–2544. doi:10.1210/jc.2004- 2232.Independent

75. Schoeller DA, Cella LK, Sinha MK, Caro JF. Entrainment of the diurnal rhythm of plasma leptin to meal timing. J Clin Invest. 1997;100: 1882–1887. doi:10.1172/JCI119717

76. Paulien Barf R, Desprez T, Meerlo P, Scheurink AJW. Increased food intake and changes in metabolic hormones in response to chronic sleep restriction alternated with short periods of sleep allowance. Am J Physiol - Regul Integr Comp Physiol. 2012;302: R112 LP-R117. doi:10.1152/ajpregu.00326.2011

77. Bodosi B, Gardi J, Hajdu I, Szentirmai E, Obal F, Krueger JM. Rhythms of ghrelin, leptin, and sleep in rats: Effects of the normal diurnal cycle, restricted feeding, and sleep deprivation. Am J Physiol - Regul Integr Comp Physiol. 2004;287. doi:10.1152/ajpregu.00294.2004

78. Koban M, Swinson KL. Chronic REM-sleep deprivation of rats elevates metabolic rate and increases UCP1 gene expression in brown adipose tissue. Am J Physiol - Endocrinol Metab. 2005;289. doi:10.1152/ajpendo.00543.2004

79. Martins PJF, Fernandes L, De Oliveira AC, Tufik S, D’Almeida V. Type of diet modulates the metabolic response to sleep deprivation in rats. Nutr Metab. 2011;8: 86. doi:10.1186/1743-7075-8-86

80. Laposky AD, Shelton J, Bass J, Dugovic C, Perrino N, Turek FW. Altered sleep regulation in leptin-deficient mice. Am J Physiol - Regul Integr Comp Physiol. 2006;290: 894–903. doi:10.1152/ajpregu.00304.2005

81. Bader R, Sarraf-Zadeh L, Peters M, Moderau N, Stocker H, Kö hler K, et al. The IGFBP7 homolog Imp-L2 promotes insulin signaling in distinct neurons of the Drosophila brain. J Cell Sci. 2013;126: 2571–2576. doi:10.1242/jcs.120261

82. Seelig JD, Jayaraman V. Feature detection and orientation tuning in the Drosophila central complex. Nature. 2013;503: 262–266. doi:10.1038/nature12601

83. Seelig JD, Jayaraman V. Neural dynamics for landmark orientation and angular path integration. Nature. 2015;521: 186–191. doi:10.1038/nature14446

84. Troup M, Yap MHW, Rohrscheib C, Grabowska MJ, Ertekin D, Randeniya R, et al. Acute control of the sleep switch in Drosophila reveals a role for gap junctions in regulating behavioral responsiveness. Elife. 2018;7: e37105. doi:10.7554/eLife.37105

85. Perkins LA, Holderbaum L, Tao R, Hu Y, Sopko R, McCall K, et al. The Transgenic RNAi Project at Harvard Medical School: Resources and Validation. Genetics. 2015;201: 843–852. doi:10.1534/genetics.115.180208

86. Shepherd D, Rokicki K, Svirskas RR, Qu L, Ngo T-TB, Laverty TR, et al. A GAL4-Driver Line Resource for Drosophila Neurobiology. Cell Rep. 2012;2: 991–1001.

87. Klapoetke NC, Murata Y, Kim SS, Pulver SR, Birdsey-Benson A, Cho YK, et al. Independent optical excitation of distinct neural populations. Nat Methods. 2014;11: 338–346. doi:10.1038/nmeth.2836

88. Kottler B, Faville R, Bridi JC, Hirth F. Inverse Control of Turning Behavior by Dopamine D1 Receptor Signaling in Columnar and Ring Neurons of the Central Complex in Drosophila. Curr Biol. 2019;29: 567–577.e6. doi:10.1016/j.cub.2019.01.017

89. Straw AD. Vision Egg: an Open-Source Library for Realtime Visual Stimulus Generation. Front Neuroinform. 2008;2: 4. doi:10.3389/neuro.11.004.2008

